# Environmental “Knees” and “Wiggles” as Strong Stabilizers of Species’ Range Limits Set by Interspecific Competition

**DOI:** 10.1101/2024.07.24.605034

**Authors:** Farshad Shirani, Benjamin G. Freeman

## Abstract

Whether interspecific competition is a major contributing factor to setting species’ range limits has been debated for a long time. Theoretical studies using evolutionary models have proposed that the interaction between interspecific competition and disruptive gene flow along an environmental gradient can halt range expansion of ecologically related species where they meet. However, the stability of such range limits has not been well addressed. We use a deterministic PDE model of adaptive range evolution over a continuous habitat to show that the range limits set by interspecific competition between two closely related species are unlikely to be evolutionarily stable if the environmental optima for fitness-related traits vary linearly in space. That is, in a (almost) linear environment without a dispersal barrier or a third (or more) related species, the range limits formed at the interface of two competing species constantly move towards the weaker species. Through extensive numerical computations, we then demonstrate that environmental nonlinearities such as “knees” and “wiggles”—wherein an isolated sharp change or a step-like change occurs in the steepness of a trait optimum—can strongly stabilize competitively formed range limits. The stabilization mechanism relies on the contrast that such nonlinearities create in the level of disruptive gene flow to the peripheral population of each species. We show that the stability of the range limits established at these nonlinearities, which are likely prevalent in nature, is robust against moderate environmental disturbances. Whether or not strong disturbances such as rapid high-amplitude changes in climate can destabilize such range limits depends on how the competitive dominance of the competing species changes across the environmental nonlinearity. Therefore, our results identify habitat regions where species ranges are fairly insensitive to climate change, and highlight the importance of measuring the competitive ability of species when predicting their response to climate change.

## Introduction

Why are species’ range limits where they are? Understanding the reasons has been a longstanding challenge in ecology [1, 2, 3, 4, 5, 6, 7, 8]. Developing such an understanding helps address a fundamental question about the future of biodiversity on the planet: where will the species’ range limits be in the near and far future? It is important, for instance, to know whether the currently contracting or expanding range of a species will eventually reach an evolutionarily stable equilibrium. This equilibrium could correspond to the species’ complete exclusion from the habitat, its full expansion over the entire available habitat, or its existence with a limited range. In the latter case, it is important to foresee where the species’ equilibrium range limits will be, whether or not they will return to the same equilibrium locations after being perturbed by changes in climate, or if they will completely be destabilized by particular climate changes. Using a mathematical model, we computationally address some of these questions for the case where range limits are formed by interspecific competitions and are stabilized by certain environmental nonlinearities.

Since Darwin, ecologists have debated the role of interspecific competition in setting species’ range limits [9, 10, 11, 12, 13]. Empiricists have used experiments and comparisons to test whether competition can limit species’ ranges [14, 15, 16, 17], while theoreticians have developed mathematical models to explore the conditions in which interspecific competition can set range limits [18, 19, 20, 21, 22, 23, 24]. The models used in these theoretical studies may or may not incorporate adaptive evolution, movements across space, and environmental heterogeneity. Non-evolutionary models often include additional factors for setting range limits, such as anti-correlated spatial profiles for species’ carrying capacity, or abrupt shifts in species’ dominance based on their resource utilization dynamics [21, 23]. However, it can be argued that, when two species are similar enough to undergo strong competition, they likely have similar resource utilization dynamics and respond similarly to abiotic environments [21]. Further, non-evolutionary models cannot address the possibility that a species may evolve to release itself from the constraining impacts of the other species. Therefore, these models cannot verify whether or not established range limits will be evolutionarily stable [23].

Among studies that consider adaptive evolution, the work of Price and Kirkpatrick [23] proposes that interspecific competition can set evolutionarily stable range limits even in the absence of disruptive effects of gene flow, and even in the case where species are competitively unequal. However, their analysis does not include species’ movement. As a result, the possibility remains that the established range limits constantly move in space due to the difference between competition strengths of the species. This means that the limits will not be evolutionarily stable. In fact, non-spatial models and models that do not incorporate movements implicitly assume there exists an external force or an internal physiological limitation on the species’ individuals which prevents them from moving to other locations. Given that even non-mobile species such as plants can sometimes disperse long distances, these assumptions are unlikely to hold for most real-world cases. The presence of dispersal-preventing forces in natural organisms is rare. Further, studying range evolution of species with restricted dispersal or movement capabilities is of less importance in understanding causes of species’ range limits.

In a seminal work incorporating species movement, Case and Taper [20] used a comprehensive evolutionary model to demonstrate that the interaction between interspecific competition and gene flow along environmental gradients can limit species’ range expansion. However, the stability of the range limits in their two-species simulations was established for the non-generic case of identical competitors, which is unlikely to ever be the case in nature. Moreover, intraspecific phenotypic variance in Case and Taper’s model [20] is constant over space and time, an assumption they describe as “tenuous”. Relaxing this assumption for a single (solitary) species results in dramatically different conclusions [25, 24], weakening confidence in the results presented by Case and Taper [20]. The main finding of Case and Taper’s influential work, that the interaction between interspecific competition and gene flow can set range limits, has been reaffirmed by Shirani and Miller [24] for the case where species’ phenotypic variance is free to evolve. However, it still remains unclear if interspecific competition can set evolutionarily stable range limits between two species in the general case of unequal competitors.

In the present work, we aim to investigate the stability of the range limits formed by interspecific competition and identify effective environmental stabilizers of such range limits. To accomplish our goals, we use the mathematical model of species’ range evolution developed by Shirani and Miller [24], an extension of Case and Taper’s model [20] that allows for the evolution of species’ phenotypic variance in space and time. Using a system of deterministic partial differential equations (PDEs), the model presents the joint evolution of the population density and the intraspecific mean and variance of a quantitative phenotypic trait for a community of competitively interacting species. To make the model better suited to our study, we further extend it to additionally incorporate Allee effects on the species’ population growth. We draw our conclusions by numerically computing the solutions of the model in different eco-evolutionary regimes.

We first show that the range limits set by interspecific competition between two species are unlikely to be evolutionary stable in a linear (or almost linear) environment (see Box 1). We then propose and analyze a reasonable stabilizing factor for the range limits: environmental nonlinearities in the form of “knees” and “wiggles” (see Box 1), the latter also briefly studied in [20] for the special case of identical competitors. These nonlinearities appear as relatively sharp changes in the steepness of the spatial profile of the environmental optimum for a fitness-related quantitative trait. Environmental wiggles and knees are likely prevalent in the real world, as nonlinear spatial changes in climate, geology, habitat structure, or biotic communities are commonly observed along environmental gradients. We show that these nonlinearities can “strongly” (see Box 2) stabilize competitively formed range limits. We further show that the established stability at the nonlinearities is fairly robust against environmental disturbances caused by climate change, and that the competitive dominance of the competing species across the nonlinearities is important for predicting possible range shifts in response to climate change.

## Materials and methods

Our work is based on a deterministic mathematical model of eco-evolutionary range dynamics in a community of competing species that disperse diffusively (randomly) in a heterogeneous environment. The model, developed by Shirani and Miller [24] as an extension of the seminal works of Kirkpatrick and Barton [26] and Case and Taper [20], has a quantitative phenotypic framework. Species’ adaptation and evolution occur based on the value of a phenotypic trait possessed by each individual in the community. The trait is assumed to be strongly correlated with individuals’ fitness. The model incorporates the effects of directional and stabilizing natural selection, which act to gradually reduce the density of the phenotypes that differ from an optimum phenotype imposed by the environment. Intra- and inter-specific competition between the species’ individuals are phenotype-dependent. Populations are assumed to be fully polymorphic. All phenotypes are assumed to be present in each population at all times, but with different frequencies. Evolution proceeds by the differential growth of phenotypes under the selective force of the frequency-dependent competition, as well as the genetic load imposed by natural selection. Phenotypic variability is assumed to be predominantly genetic, and genetic variation is assumed to be predominantly additive. That is, H^2^ ≈ h^2^ ≈ 1, where H^2^ and h^2^ denote broad- and narrow-sense heritability, respectively.

### Box 1

**Ecological and evolutionary terminology**

**Environmental gradients**

In general, an *environmental gradient* along a geographic space refers to a spatial change in the environmental conditions that affect species’ fitness and performance. Such changes in environmental factors can cause, for example, a gradient in species’ productivity which can be modeled by spatial changes in the carrying capacity of the environment. Environmental conditions such as temperature, humidity, light density, and resource quality can also cause gradients in the most suitable (optimum) phenotypes. We say an environment is *linear* if the effects of all environmental conditions on the fitness (growth rate) of the species vary linearly in space, that is, with constant gradient. Throughout the present work, an environmental gradient always refers to a selective gradient that the environment imposes on the individuals by setting the optimum value of a fitness-related quantitative trait (phenotype). Therefore, the environment is linear if the trait optimum changes linearly in space. Other environment-dependent parameters are assumed to be constant in space. However, our results remain qualitatively valid if changes in such factors occur linearly.

**Environmental knees**

By *environmental knee* we refer to a point along an environmental gradient in trait optimum at which the steepness of the gradient changes sharply; see the black line in Figure 3b.

**Environmental wiggles**

**Table 1.**
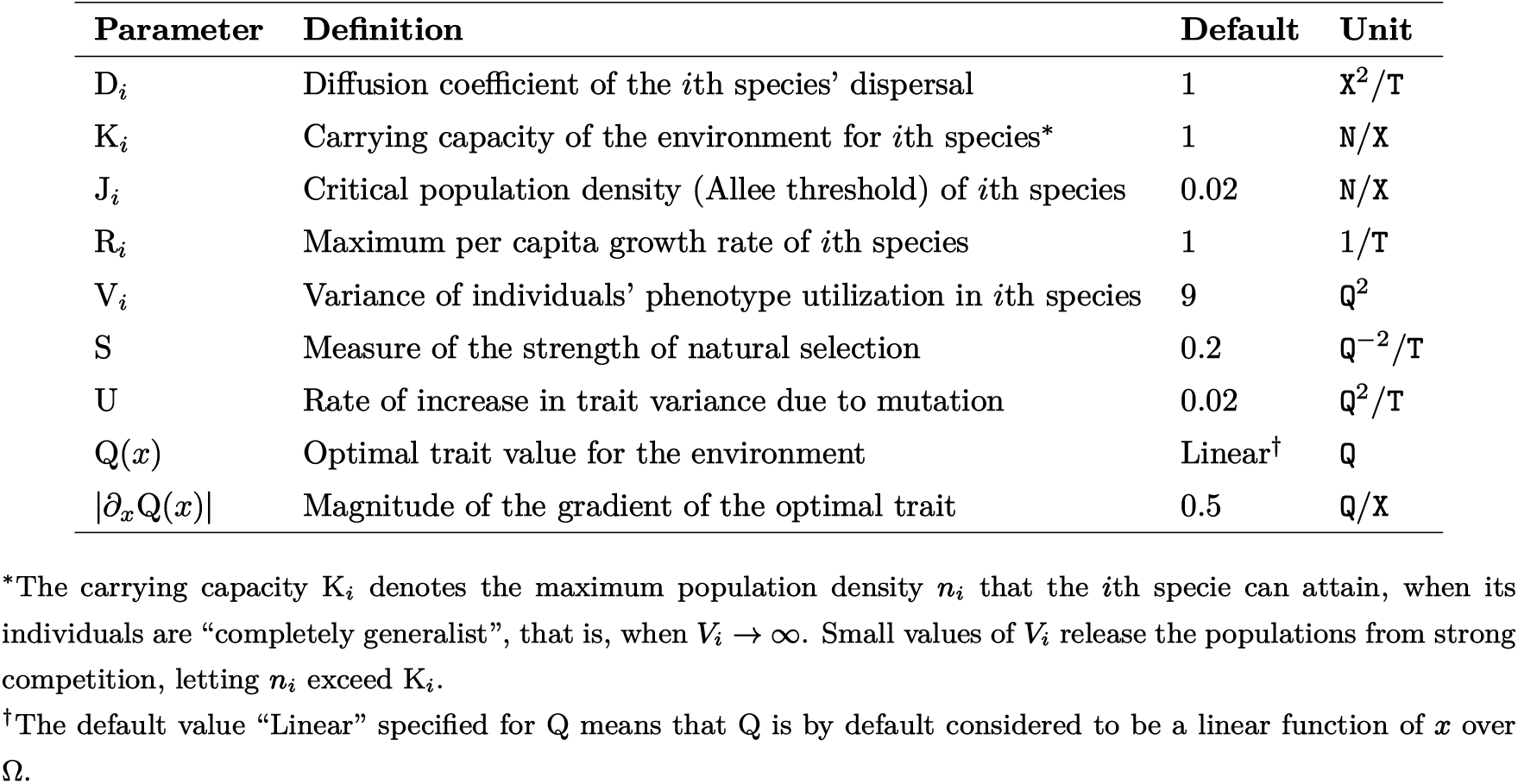
Definition and default values of the parameters of the model (5)–(7). The default values are used in the simulations presented in the Results section, unless otherwise is stated.

By *environmental wiggle* we refer to a step-like nonlinearity in the environmental trait optimum, across which the steepness of the gradient changes sharply from an initially moderate value to a significantly higher value, and then back to a moderate value; see the black line in Figure 3d. This means that, outside a wiggle the gradient is moderate/shallow, whereas inside the wiggle it is steep. A wiggle is therefore composed of two adjacent knees, with opposite directions of changes in the gradient steepness. We further require the spatial span of the nonlinearity to be relatively short; approximately shorter than ten times the mean dispersal distance of the individuals in one generation time. Exceedingly wide wiggles are better understood as two separate knees. We refer to the geographic region within a wiggle (i.e., between the two knees) as *transition zone* of the wiggle, and to the steepness of the environmental gradient over the transition zone as *steepness* of the wiggle.

**Fitness loads**

In general, we refer to any factor that causes a reduction in the fitness (growth rate) of individuals, or the mean fitness of populations, as a *fitness load*. Depending on the nature of the factor and the fitness component it affects, we may further specify it as a *migration load* (caused by random gene flow), *genetic/phenotypic load* (caused, for example, by natural selection), *competition load*, or *genetic drift load*.

**Phenotype utilization distributions**

The *phenotype utilization distribution* (curve) of an individual with phenotype *p* is a probability density function, which we denote by *ψ*_*p*_. When evaluated at a phenotype value 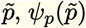 gives the probability density that an individual with phenotype *p* will utilize a unit of resource that is most favorable for (i.e., is expected to be mostly utilized by) an individual with phenotype 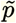. Assuming that the environmental resources vary continuously along a resource axis, and that there exists a one-to-one mapping between the resource values and phenotype values, the phenotype utilization curves are computed based on resource utilization curves through identifying the resource axis by the phenotype axis [18, Eq. (24.5)]. See Appendix A.2 and Section 3.2 in [24] for further details.

We additionally incorporate Allee effects into the model. The population growth rate in Shirani and Miller’s model [24], as well as in Case and Taper’s model [20], is assumed to be purely logistic. This means that the growth rate of a population becomes maximum when its density goes to zero. As a result, any infinitesimal population of a competitively stronger species that initially coexists with the weaker species, or gradually leaks to the region where the weaker species is established, can grow to significantly large densities and likely exclude the weaker species from the habitat. This is a biologically unrealistic situation. Populations of very low density are subject to Allee effect and a strong load of random genetic drift such that they are unlikely to survive, especially in the presence of an abundant competitor [27, 28]. The eco-evolutionary regimes studied in [24] do not suffer from this unrealistic situation. However, our analyses in the present work require simulations (computations) over very long evolutionary time horizons, to ensure the stability of the interface formed between two competing species at environmental knees and wiggles. The Allee effects that we add to the model, as described below, help prevent the infinitesimal populations to unrealistically disturb the results. In the presence of Allee effects, the populations whose density is below a critical (small) density will have a negative growth rate and will fail to survive.

We obtain our results by numerically solving the equations of the model for two ecologically related species, one of them being (generally) competitively stronger than the other, living along a one-dimensional geographic region (habitat). We use the same numerical method as used in [24] to solve the equations, with slight modifications. Further details of the numerical scheme and technical challenges in computing the solutions are provided in Appendix B. We model an environmental knee by sharply changing the steepness of the environmental gradient (in the trait optimum) at a given geographic point, and an environmental wiggle by making a step-like change in the trait optimum. We analyze the evolution and stability of the species’ competitively formed range limits as they enter and get stabilized in an environmental knee or wiggle. We test the robustness of the stability of the range limits against moderate and strong climate-warming disturbances.

It is important to note that our numerical simulation-based analyses cannot serve as mathematical proofs of stability of the range limits. A rigorous mathematical analysis of the stability of the range limits in the PDE model that we use, in an infinite-dimensional dynamical systems framework, is quite challenging and is beyond the scope of this work. In the results we present, the coevolutionary mechanisms of range stabilization that we describe in words are the main arguments that convince us of the stability (and its type) of the range limits formed at environmental knees and wiggles. Our numerical simulations are mainly purposed to demonstrate and support our verbal descriptions.

Before presenting the results, we describe the essential components of the model governing the underlying dynamics of the species’ range evolution. We present the formulations for a reduced version of Shirani and Miller’s model [24] for two interacting species in a one-dimensional geographic space. We refer the reader to their work for further details of the model for multiple species in higher dimensional spaces. Our new addition of the Allee effects slightly changes the formulations of the final equations of the model as presented in [24]. We provide the complete set of model equations in Appendix A, along with its reduction to a single-species case. Below, we describe the basic components.

### Model Components

We describe the model for adaptive range evolution of two species in a one-dimensional habitat, over an evolution time horizon from *t* = 0 to *t* = *T >* 0. We denote the habitat by an open interval Ω = (*a, b*) ⊂ ℝ. At every location *x* ∈ Ω and time *t* ∈ [0, *T*], the *i*th species’ range dynamics is governed by three coupled equations that control the joint evolution of the population density *n*_*i*_(*x, t*), intraspecific trait mean *q*_*i*_(*x, t*), and intraspecific trait variance *v*_*i*_(*x, t*), where *i* = 1, 2. These main equations of the model are given as equations (5)–(7) in Appendix A. The description of the basic contributing components of the model are as follows.

Let *ϕ*_*i*_(*x, t, p*) denote the relative frequency of phenotype value *p* ∈ ℝ within the *i*th species’ population. Then *n*_*i*_(*x, t*)*ϕ*_*i*_(*x, t, p*) gives the population density of individuals with phenotype *p* in the *i*th population. The model assumes that changes in *n*_*i*_(*x, t*)*ϕ*_*i*_(*x, t, p*) over a small time interval of length *τ* → 0 results from the contribution of three major factors, as:

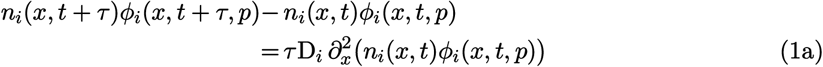

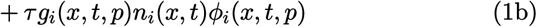

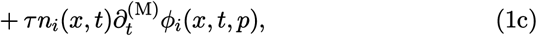

where 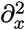 denotes the second partial derivative with respect to *x*. The first contributing factor (1a) represents the diffusive (random) dispersal of individuals to and from neighboring locations, where D_*i*_ denotes the diffusion coefficient; see Table 1. The second contribution (1b) is due to the intrinsic growth of the individuals with phenotype *p* in the *i*th population, the rate of which is denoted by *g*_*i*_(*x, t, p*). Finally, the third factor (1c) represents mutational changes in the relative frequency of the phenotype *p*, occurring at a rate denoted by 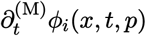. The probability of mutational changes from a phenotype *p* to another phenotype *p*′ is assumed to be dependent on the difference *δp* = *p* − *p*′ between the phenotypes. Letting *ν*(*δp*) denote the probability density of the occurrence of such mutational changes, the model further assumes that *ν* follows a distribution with zero mean and constant variance. Based on these assumptions, the formulation used in the model for 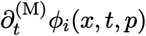 is given in [24, Section A.3].

The key factor that differentiates the density of individuals with different phenotypes is their intrinsic growth rate, which is modeled as,

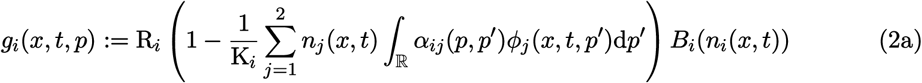

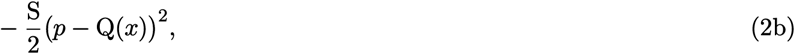

where S denotes the strength of stabilizing selection, Q(*x*) denotes the environment’s optimal trait value at location *x*, R_*i*_ denotes the maximum per capita growth rate of the *i*th species, and K_*i*_ denotes the carrying capacity of the environment for the *i*th species when the species’ individuals are completely generalist; see also Table 1. The rest of the terms are described below. Note that, in writing (2a), it is essentially assumed that the reproduction rate of the individuals with phenotype *p* follows a logistic growth (with Allee effect), with R_*i*_ and K_*i*_ being independent of their phenotype.

The convolution term in (2a) models the effects of phenotypic competition between the individuals. Specifically, the competition kernel *α*_*ij*_(*p, p*′) denotes the strength of the per capita effects of individuals with phenotype *p*′ in the *j*th species on the frequency of individuals with phenotype *p* in the *i*th species. To specify this competition kernel, the model uses the MacArthur-Levins overlap [29] between *phenotype utilization curves* of the individuals (see Box 1), assuming that the environmental resources vary continuously along a resource axis. The phenotype utilization distribution (curve) associated with individuals with phenotype *p* in the *i*th population is assumed to be normal, with mean *p* and constant utilization variance V_*i*_. The overlap between the phenotype utilization curves gives the competition kernel as

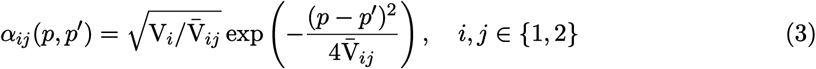

where 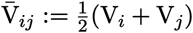. The details are given in [24, Sections A.2 and A.4].

Before proceeding to define the remaining terms in (2), it is important to clarify an ambiguity in the definition of K_*i*_ that is present due to the phenotype-dependence of competition in the model. As stated above, K_*i*_ denotes the maximum population density of the *i*th species when the species’ individuals are “completely generalist”, that is, when V_*i*_ → ∞, *i* = 1, 2. This is because *α*_*ij*_(*p, p*′) → 1 in (3) when 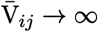, regardless of the values of *p* and *p*′. Setting *α*_*ij*_(*p, p*′) = 1 in (2) verifies that completely generalist species with density *n*_*i*_ *>* K_*i*_ will have negative intrinsic growth. However, when individuals are specialized in utilizing resources, they become partially released from competition, meaning that *α*_*ij*_(*p, p*′) *<* 1. In this case, *g*_*i*_ can take positive values for some *n*_*i*_ *>* K_*i*_. That is, specialized species can grow to population densities greater than K_*i*_. For example, see the curves of range expansion wave amplitudes (maximum population density) given in [30, Fig. 2b]. Therefore, even though K_*i*_ is phenotype-independent, the carrying capacity of species— defined generally as species’ maximum population density—depends both on the distribution of phenotypes and individuals’ phenotype-based resource utilization. Unless explicitly specified as K_*i*_, in our references to carrying capacity in the rest of this paper we always mean populations’ maximum attainable density.

The nonlinear function *B*_*i*_ of the species’ population density in (2a) presents the Allee effect. We assume that, due to the Allee effect, a population whose density is below a critical density, called the *Allee threshold*, will have a negative intrinsic growth rate. We denote the critical density for the *i*th species with J_*i*_. When the population’s density increases above the critical density, the population’s intrinsic growth rate increases sharply to the maximum rate R_*i*_. Further increases in the density results in a gradual decrease in the growth rate, eventually to zero when the density reaches the carrying capacity. For this, we define *B*_*i*_ as

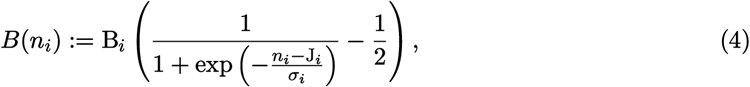

where *σ*_*i*_ is a parameter for adjusting the sharpness of the growth rate increase to its maximum, and B_*i*_ is a parameter for adjusting the maximum growth rate (to be R_*i*_). For both species throughout our study, we set J_*i*_ = 0.02 N*/*X, *σ*_*i*_ = 0.05 N*/*X, and B_*i*_ = 2.6. The resulting profile of the intrinsic growth rate (2) for a fully adapted solitary population with completely generalist individuals is shown in Figure S1 of the supplementary document.

Finally, the quadratic term (2b) in the intrinsic growth function imposes directional and stabilizing forces of natural selection on individuals with phenotype *p*. It penalizes phenotypes that are far from the environmental optimal phenotype Q(*x*), thereby facilitating adaptation to new environments. In our results, we model environmental wiggles by inserting short-range step-like nonlinearities in the otherwise linear profile of Q.

The phenotype density evolution equation (1), along with the definition of its components as described above, is used to derive the final equations of the model for the evolution of *n*_*i*_, *q*_*i*_, and *v*_*i*_, as given in equations (5)–(7) in Appendix A. The Allee effect term *B*_*i*_(*n*_*i*_) that we have included in (2) does not depend on the phenotype *p*. As a result, following the derivations provided in [24] simply yields a replacement of R_*i*_ with *B*_*i*_(*n*_*i*_)R_*i*_ in the final equations derived in [24]. We note that, the master equation (1) is specified at the phenotypic level. Therefore, the derivation of the equations in [24] is performed entirely at the phenotypic level. The derivation relies on a key assumption, inherited from the preceding foundational models [20, 26], that phenotype values in each species are normally distributed at every occupied habitat location for all time. This assumption allows for exact moment closure in deriving the equations for trait means (by computing first moment of phenotypes) and trait variances (by computing the second moment of phenotypes). See Appendix A.3 for a discussion on the validity of this assumption in our study. As stated above, in deriving the equations of trait variance, it is also implicitly assumed in [24] that phenotypic variation is predominantly caused by additive genetic variation (H^2^ ≈ h^2^ ≈ 1).

The definitions of the model parameters described above are given in Table 1, along with their default values used in our simulations. The choice of units for the physical quantities of the model is an important factor in suggesting biologically reasonable values for model parameters based on available estimates in the literature. Due to the complexity of the equations, analytical studies of the behavior of the model in different evolutionary regimes will be rather impractical. Numerical studies of the model, as we perform in this work, will then require carefully chosen parameter values such that the resulting predictions are biologically reasonable. An extensive discussion is made in

[24] on the choice of units and reasonable ranges of parameter values. Below, we only describe the choices of units as needed for understanding the values given in Table 1, and provide some examples on how empirical measurements would be presented based on such units.

### Parameter Units

To specify units for the physical quantities of the model, one of the species in the community is first chosen as a *representative species*, for example, the one which is best adapted to the environment or has the widest niche. The units are then specified based on the measurements made on the representative species. Specifically, the unit of time, denoted by `T`, is set to be equal to the mean generation time of the representative species. The unit of space, denoted by `X`, is chosen such that the diffusion coefficient D of the representative population becomes unity. That is, 1 `X` is set to be equal to the root mean square of the dispersal distance of the population in 1 `T`, divided by 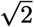.

Having set the unit of space, the unit of measurement for population abundances, `N`, is then chosen such that 1 `N` is equal to the carrying capacity of the environment for 1 `X` unit of habitat length. Note that this results in the default value of K_*i*_ for the representative population to become unity, provided the individuals of the population are sufficiently generalist. Finally, letting `Q` denote the unit of measurement for the quantitative trait, 1 `Q` is set to be equal to one standard deviation of the trait values at the core of the representative population.

As shown in [31, Sect. 3.2], the strength of natural selection S can be estimated as S ≈ − *γ* + *β*^2^, where *γ* denotes the standardized stabilizing selection gradient and *β* denotes the standardized directional selection gradient [32, 33, 34]. Estimates of these standardized selection gradients are available in the literature, see for example [33, 35, 36]. These estimates can be directly used to provide an estimate for S in our model, as they are obtained based on one generation (our unit of time) of selection for a trait measured in standard deviation units (our unit of trait). Estimates for the mutational rate of increase in trait variance are available in the literature, either relative to environmental variance or relative to standing genetic variance [37]. Since we choose standard deviation of the trait in the representative population as the unit of trait, the standing genetic variance can be approximated to be unity. Since it is also implicitly assumed in our model that phenotypic variation is almost equal to genetic variation (H^2^ ≈ 1), the available estimates in [37] can be used directly as approximate estimates for U in the model we use; see [24] for further discussion. Estimates for the variance of phenotype utilization distribution V_*i*_ can be obtained by computing the within-phenotype component of species’ niche breadth, as defined in trait-based niche quantification approaches [38, 39, 24]. Based on our choices of unit for trait mean, the estimates are expected to be greater than 1, since the local representative population might not yet have filled its entire possible niche.

We use a toy examples to illustrate how parameter values measured in arbitrary units are converted to values based on our choices of units. Consider a species of birds whose individuals’ wing length (trait) increases as the partial pressure of oxygen decreases across an elevational gradient. Suppose a well-adapted representative local population of this species has a trait distribution with standard deviation equal to 5 mm. Suppose further that the environmental optimum for the trait increases linearly across the habitat, with a slope equal to 2 mm*/*km. The individuals are generalist, and the carrying capacity of the environment is equal to 4000 individuals per kilometer. Note that, for simplicity, we assume the habitat is one-dimensional. The maximum per capita intrinsic growth rate of the individuals is 0.3 per year. The mean generation time of the species is 3.5 years and its mean dispersal distance is 0.7 kilometer per year. An estimate for individuals’ phenotype utilization variance obtained based on trait-based niche quantification is equal to 100 mm^2^. Estimates for the standardized directional and stabilizing selection are *β* = 0.2 and *γ* = −0.15, based on one generation of selection and for wing lengths measured in standard deviation units. For this example species, we compute the parameter values based on our choices of units, as follows.

The unit of time is set equal to the mean generation time, 1 `T` = 3.5 years. The mean dispersal distance per generation is equal to 0 7 3 5 = 2 45 kilometers. Therefore, we set the unit of space as 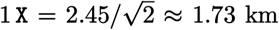. The carrying capacity for one unit of habitat length is then 4000 × 1.73 = 6920 individuals. We set the unit of population abundance as 1 N = 6920 individuals. Finally, we set the standard deviation unit for trait, that is 1 Q = 5 mm. Based on these units, it follows immediately that D = 1 `X`^2^*/*`T`, and K = 1 `N`*/*`X`. The maximum intrinsic growth rate is R = 0.3 3.5 = 1.05 `T`^−1^. The phenotype utilization variance is V = 100*/*5^2^ = 4 `Q`^2^. The strength of selection is S ≈ −*γ* + *β*^2^ = 0.15 + 0.2^2^ = 0.19 Q^−2^*/*T. Finally, the slope of the environmental gradient is computed as 2 × 1.73 = 3.46 mm*/*`X`, which gives the slope *∂*_*x*_Q = 3.46*/*5 ≈ 0.7 `Q`/`X` based on our units. The estimates in [24] suggest that this environmental gradient is quite steep.

## Results

We investigate how the interaction between interspecific competition and gene flow leads to formation of range limits between two competing species, and how the limits are stabilized by environmental knees and wiggles. For this, we numerically solve the equations of the model (5)–(7) given in Appendix A.1, with their nonlinear terms given by (8)–(17). Since the maladaptive effects of random dispersal (gene flow) plays an important role in the evolutionary processes we describe in this work, we first demonstrate such effects using a reduction of the model to a single-species case (Appendix A.2). We then demonstrate the basic process of character displacement and formation of range limits between two species by parameterizing the two species with identical ecological and evolutionary parameter values; equal to the default values given in Table 1. Following these preliminary analyses, we then make specific changes in the maximum growth rate of each species to further investigate the generic case of competitively unequal species.

We perform our simulations in the one-dimensional habitat Ω = (0 `X`, 50 `X`) ⊂ ℝ with the reflecting boundary conditions (18). We set the spatial profile of the trait optimum Q to be decreasing along the habitat. This simplified habitat can be conceptualized to represent, for example, the living environment of two species of montane birds over an elevation-dependent temperature gradient, with Q representing a temperature-correlated phenotypic trait (for example, clutch size is a trait in birds which decreases with elevation [40]). The reflecting boundary condition at *x* = 50 can then be equivalent to symmetrically (reflectively) expanding the habitat at the mountain top. Our choice of the default value R_*i*_ = 1 `T`^−1^ in Table 1 is also purposed to agree with a typical value of the maximum growth rate per generation for birds [41, 42, 24].

In all simulations, except the first single-species simulation, we initially introduce the two populations allopatrically, considering the 1st population as the *downslope* (at lower elevations) and the 2nd population as the *upslope* species. We set the initial population densities at *t* = 0 `T` as *n*(*x*, 0) = 0.5*ϱ*((*x* − *c*_*i*_)*/*2), where *c*_*i*_ denotes the center of the *i*th population and *ϱ* is a bump function with support [−1, 1]; see also Appendix B. We assume that the initial populations are perfectly adapted to their environment at their center, that is, *q*_*i*_(*c*_*i*_, 0) = Q(*c*_*i*_). However, we assume fairly poor initial adaptation at other habitat locations by letting the slope of the initial trait means vary as *∂*_*x*_*q*_*i*_(*x*, 0) = 0.1 *∂*_*x*_Q(*x*), for all *x* ∈ Ω. Finally, we assume that the initial populations have a constant trait variance of *v*_*i*_(*x*, 0) = 1 Q^2^. For the single-species case, we consider the same simulation layout but without introducing the 2nd species.

### Range Expansion Dynamics and Maladaptive Effects of Random Gene Flow

The range limits stabilization mechanism that we described in the present work relies on maladaptive gene flow from core to peripheral populations, which is created by random dispersal of individuals. To better understand the stabilization mechanism, we first demonstrate the range expansion dynamics of a solitary population in a linear environment and explain how it is affected by gene flow, selection, and intraspecific competition. For this, we solve the equations of the model for a single species, which take the simpler form given in Appendix A.2. The simulation results are shown in Figure 1.

**Figure 1:**
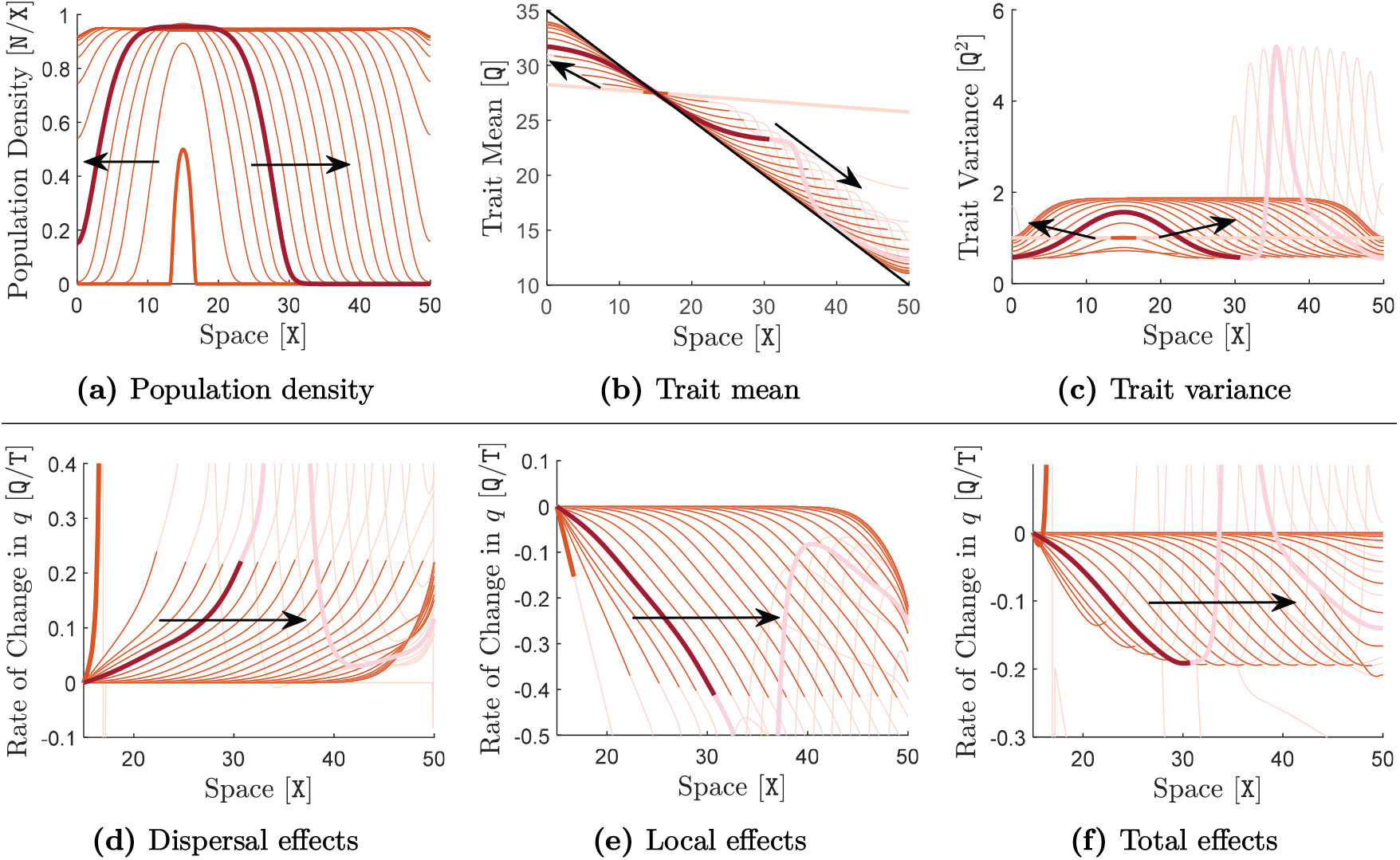
Range evolution of a solitary species and the effects of random gene flow, selection, and competition on population adaptation. The parameters of the species are set equal to the default values given in Table 1. The trait optimum Q, shown by the black line in (b), changes linearly with a moderately steep gradient (slope) of *∂*_*x*_Q = −0.5 `Q`*/*`X`. Curves of the species’ population density, as its range evolves in time, are shown in (a). The corresponding curves of species’ trait mean and trait variance are shown in (b) and (c), respectively. The contributions of dispersal (gene flow) as well as the local contributions of selection and competition to the rate of change of trait mean *∂*_*t*_*q* is shown in (d) and (e), respectively. The curves are shown only for the right-half of the species range (15 `X`, 50 `X`). Curves in (d) are computed using the term D*∂*^2^*q* + 2D*∂*_*x*_(log *n*)*∂*_*x*_*q* in (20), and curves in (e) are computed using the term −S(*q* − Q)*v*. The total contribution to *∂*_*t*_*q*, shown in (f), is computed as the sum of these two terms, that is, the sun of the curves shown in (d) and (e). In all graphs, curves are shown at every 4 `T` for a simulation time horizon of *T* = 200 `T`. The parts of the curves which lie outside the species’ range (*n <* 0.02) are made transparent, as the values of trait mean and trait variance over these regions are not biologically meaningful. The thick orange curves indicate the initial curves at *t* = 0 T, and arrows show the direction of evolution in time. In each graph, a sample curve at *t* = 20 T is highlighted in red.

As the initially introduced population establishes itself in the habitat, it gradually adapts to new environments under the force of natural selection (Figure 1b) and expands its range in the form of traveling waves (Figure 1a). At the core of the population, the density of the population grows to the environment’s maximum capacity and its trait mean converges to the environmental trait optimum. The population’s density declines near the edge of its range, which creates an asymmetry in gene flow from the populous core of the population to its edge. Due to the presence of a gradient in trait optimum, the phenotypes that are adapted to central regions of the range will be maladapted at peripheral regions. Therefore, the asymmetric gene flow decreases the mean fitness of the population at its range margins. As a result, population density decreases further near the range edge, which then results in a further increase in the asymmetry of gene flow. This positive feedback loop between population density and gene flow slows down the range expansion of the population. The steeper the environmental gradient, the stronger the maladaptive effects of gene flow and the slower the range expansion speed; see the curves of range expansion speed in [24, Fig. 4] and [30, Fig. 2] for a similar simulation.

**Figure 2:**
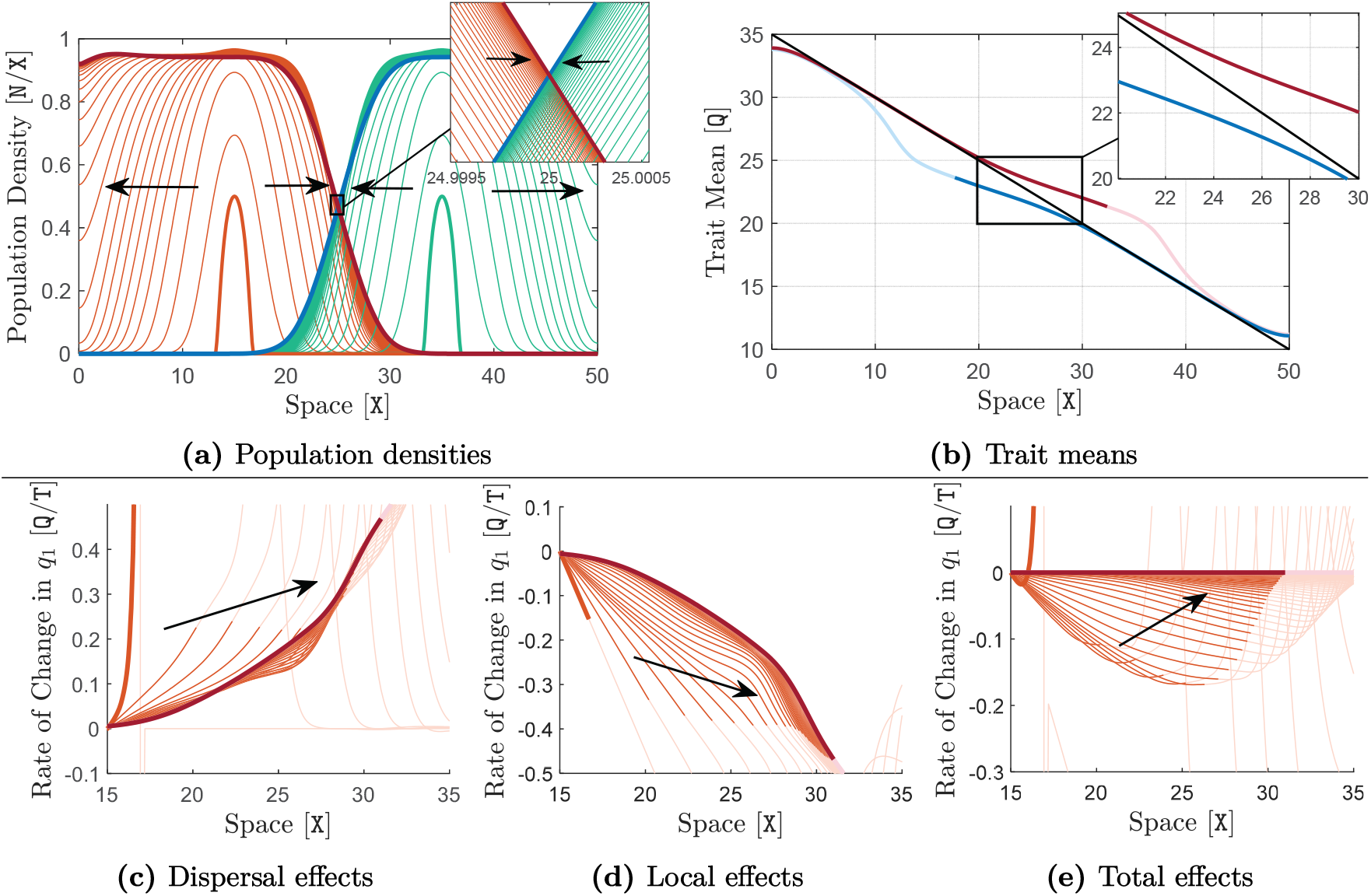
Evolution of range limits, region of sympatric coexistence, and character displacement for two competitively identical species. Both species have the same parameter values equal to the default values given in Table 1. The trait optimum Q is linear and decreasing, shown by the black line in (b), with a moderately steep gradient of *∂*_*x*_Q = −0.5 `Q`*/*`X`. Curves of the species’ population density are shown in (a). The final curves of the species’ trait mean obtained at the end of the simulation are shown in (b). The contribution of dispersal (gene flow) to the rate of change of trait mean *∂*_*t*_*q*_1_ is shown in (c). The local contributions of selection and competition to *∂*_*t*_*q*_1_ are shown in (d). The curves are shown only for the right-half of the 1st species’ range, (15 `X`, 50 `X`). Curves in (c) are computed using the term 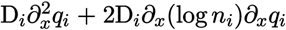 in (6), with *i* = 1. Curves in (d) are computed using the term *H*_*i*_ in (6), with *i* = 1. The total contribution to *∂*_*t*_*q*_1_, shown in (e), is computed as the sum of the curves shown in (c) and (d). For the downslope (left) species, curves are shown in orange, thick orange curves indicate the initial curves at *t* = 0 `T`, and the final curves at the end of the simulation are highlighted in red. For the upslope (right) species, curves are shown in green, thick green curves indicate the initial curves, and the final curves are highlighted in blue. Arrows show the direction of evolution in time. In (a) and (c)–(e), curves are shown at every 2 T for a simulation time horizon of *T* = 200 T. The parts of the curves which lie outside the species’ range (*n*_*i*_ *<* 0.02) are made transparent. The ranges converge to an equilibrium state, with a limited region of sympatry formed in the middle of the habitat. Graph (b) shows that the species evolve character displacement over the region of sympatry.

**Figure 3:**
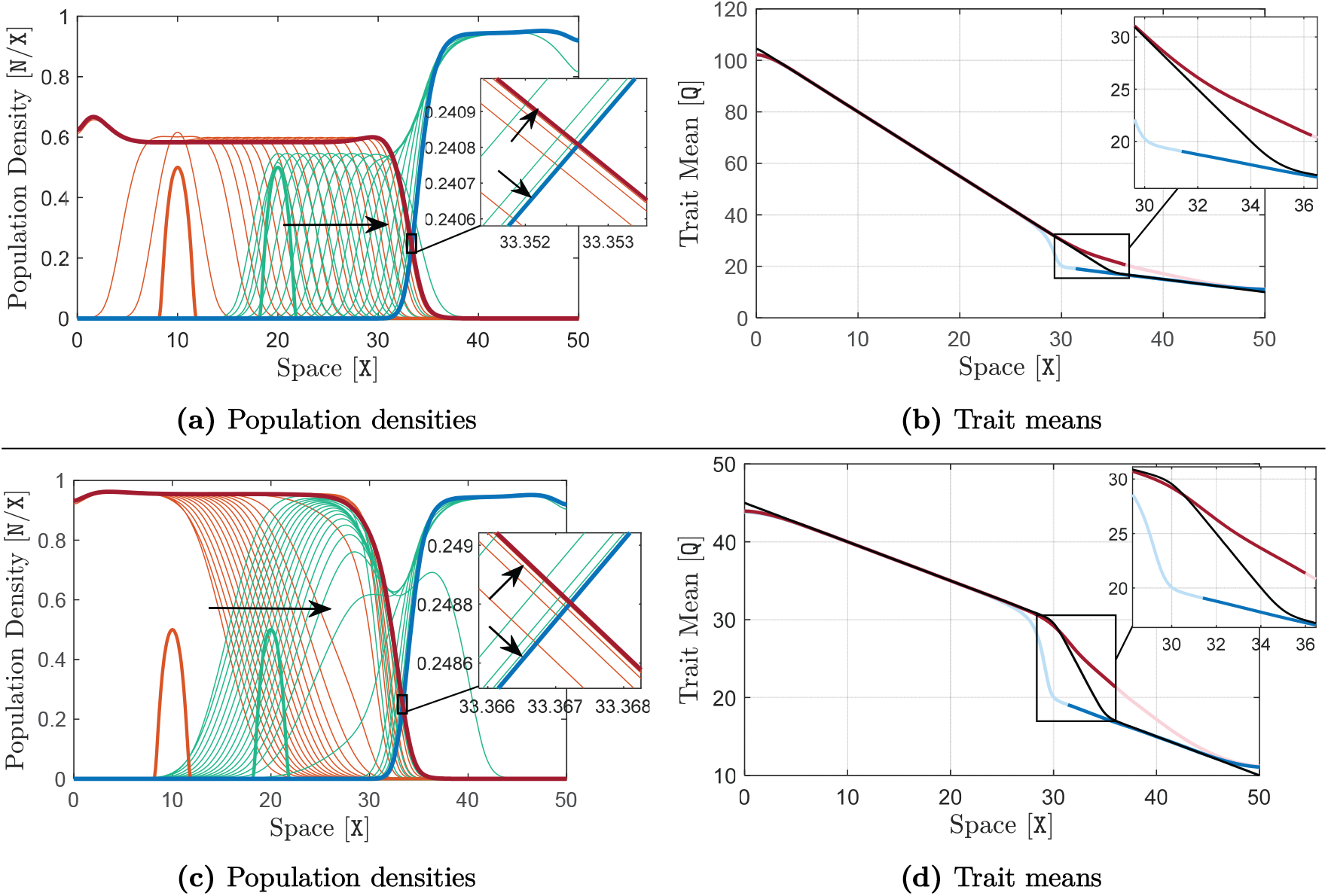
Formation of stable range limits at environmental knees and wiggles. Here, R_1_ = 1.1 `T`^−1^, R_2_ = 1 `T`^−1^, and the rest of the parameters take their typical values given in Table 1. The results shown in (a) and (b) in the upper panel are associated with a habitat with an environmental knee located at *x* = 35 `X`, at which the slope of spatial variations in the trait optimum changes sharply from − 2.5 `Q`*/*`X` to − 0.5 `Q`*/*`X`. The results shown in (c) and (d) in the lower panel are associated with a habitat with an environmental wiggle, at which the slope of trait optimum switches sharply from − 0.5 Q*/*X to − 2.5 Q*/*X between the two knees of the wiggle located at *x* = 30 `X` and *x* = 35 `X`. The simulations are performed for a time horizon of *T* = 1500 T and curves of population density are shown in (a) and (c) at every 30 `T`. The same description as given in Figure 2 holds for the curve colors and arrows. The curves highlighted in red and blue are associated with the equilibrium state of the populations, reached (approximately) at the end of the simulations (*t* = 1500 T). The highlighted equilibrium curves in (a) and (c) represent the evolutionarily stable range limits formed at the environmental knee and wiggle, respectively. The corresponding curves of equilibrium trait means are shown in (b) and (d), where the black lines show the environmental trait optimum Q. The equilibrium curves of trait mean are made transparent over the regions where population densities are approximately zero.

**Figure 4:**
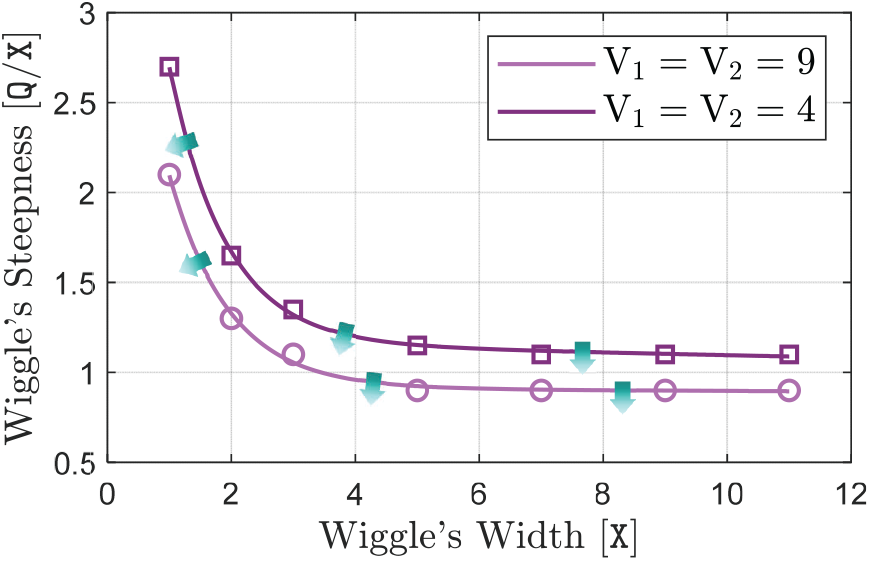
Critical wiggle parameters for stabilizing the range limits. To identify the critical wiggle parameters for stability of the range limits, the simulation associated with the lower panel of Figure 3 is repeated here for different values of steepness and width of the wiggle. The slope of the environmental gradient outside the wiggle is kept fixed at the same value −0.5 `Q`/`X` as used in Figure 3. To find the critical wiggle parameters, approximately, different values are chosen for the wiggle’s width within a reasonably wide range. A step-wise search (step-wise changes of step size 0.05 `Q`/`X` in the steepness) is then performed at each width value to find the critical value of the wiggle’s steepness, below which the wiggle fails to stabilize the range limits. To do the searches, the simulation is run at each step (i.e., for each (width, steepness) pair) for a period of *T* = 5000 T and convergence to an equilibrium is checked at the end of the simulation. The critical parameter values obtained through the searches are marked in the graph. The critical values marked by circles are associated with simulations with the strong level of competition (V_1_ = V_2_ = 9 Q^2^) used throughout the paper. The critical values marked by squares are associated with a more moderate level of competition (V_1_ = V_2_ = 4 Q^2^). The curves connecting the points are computed by interpolation. Wiggles that are shallower or narrower than the critical values marked in the graph (lie below the curves) cannot stabilize the range limits (i.e., no convergence to an equilibrium is verified at the end of the simulation). Arrows indicate transition from stable (dark) to unstable (light) range limits as the wiggle parameters change across the curves.

To better illustrate the maladaptive effects of random dispersal, the direct effects of dispersal on rate of change of trait mean *∂*_*t*_*q* is shown in Figure 1d, computed using the term 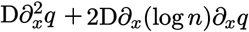 in (20). The curves in Figure 1d are shown for the right-half of the species’s range, where its trait mean initially lies above the trait optimum at *t* = 0 T. Therefore, adaptation occurs when the trait mean decreases to the trait optimum, that is when *∂*_*t*_*q* takes negative values. Positive values of *∂*_*t*_*q* will then imply maladaptive effects. As we see in Figure 1d, the effects of random dispersal that brings phenotypes from the core to the edge are maladaptive at range margins.

The gene flow along the gradient will, on the other hand, increase genetic variation in local populations; the steeper the gradient the higher the level of genetic variation [30, Figure 2a]. In the models that assume constant phenotypic variance, the swamping effects of asymmetric gene flow on adaptation at the range edge will become so strong at steep gradients that range expansion will halt [26]. However, when we allow trait variance to evolve (Figure 1c), the inflation in phenotypic variation caused by gene flow facilitates adaptation by natural selection. This is because increased trait variation provides more opportunities for natural selection to operate, allowing for local adaptation and range expansion even at exceedingly steep gradients. Figures 1d–1f illustrate the overall mechanism of adaptation at range margins. The term −S(*q* − Q)*v* in (20), shown in Figure 1e, represents the local adaptation by selection acting on the genetic variation created by gene flow, mutation, and intraspecific competition. We note that in relatively steep gradients such as that simulated in Figure 1, the effects of mutation and intraspecific competition on creating genetic variation are rather insignificant compared with the dominant effects of gene flow. As Figures 1d–1f show, the local adaptation caused by selection is strong enough to overcome the maladaptive effects of random dispersal, allowing for adaptive range expansion to new environments. Note that negative values in Figures 1d–1f imply adaptive effects, because the trait mean on the right-half of the species’ range is initially above the trait optimum.

The inflating effects of random gene flow on trait variation and thereby increasing the population’s adaptive potential creates ambiguities on our general reference to “maladaptive” effects of gene flow, which we need to clarify. The results shown in Figure 1 were obtained for a moderately steep gradient of *∂*_*x*_Q = −0.5 `Q`*/*`X`. To clarify the effects of gene flow, we repeat the same simulation but with a much shallower gradient of *∂*_*x*_Q = −0.1 `Q`*/*`X` as well as a much steeper gradient of *∂*_*x*_Q = −2.5 `Q`*/*`X`. The results are shown in supplementary Figure S2. Comparing Figures 1d, S2a, and S2d confirms that the disruptive effects of core-to-edge gene flow, which deviate the trait mean from trait optimum, increase as the level of gene flow increases with steepness of the environment. In contrast, Figures 1e, S2b, and S2e confirm that the adaptive potential of the peripheral population increases with the steepness of the environment. This is because trait variation increases as the population disperses and adapts along steeper gradients, mainly due to the term 2D |*∂*_*x*_*q*| ^2^ in (21). The sum of these two contrasting effects gives the total rate of change in trait mean *∂*_*t*_*q*, shown in Figures 1f, S2c and S2f, which changes in a more complicated way with increases in environmental gradient.

Noting that more negative values of *∂*_*t*_*q* in the results presented here imply faster adaptation, Figures 1f, S2c and S2f show that adaptation rate at range margins is higher when the environment is moderately steep. This could imply that the overall effects of intermediate levels of gene flow is adaptive at range margins, and hence could facilitate range expansion compared with low or very high levels of gene flow. However, examining the range expansion speeds in Figures 1 and S2, as well as the speeds shown in [24, Fig. 4] and [30, Fig. 2a] contradicts this conjecture. Range expansion speed decreases monotonically as the steepness of the environmental gradient and hence the level of gene flow increases. This means that, even though the maladapted phenotypes that are brought to the range periphery from the range core also create an adaptive potential to mitigate their maladaptive effects, the migration load they impose on the population fitness (deviation of the trait mean from the trait optimum [43]) is always too strong to be fully compensated by the effects of increased adaptive potential. To further elaborate on this, we repeat the simulation associated with Figure 1 but with an initial population that is perfectly adapted to the trait optimum, *q* = Q at *t* = 0 T. The results are shown in Figure S3. If the overall effects of gene flow were adaptive, this initial population would remain perfectly adapted for all time. Instead, we see that gene flow quickly deviates the trait mean of the peripheral population from the optimum, and adaptation and range expansion proceeds almost the same as in Figure 1.

It is worth pointing out that the overall adaptation at range expansion wavefronts, as in Figures 1f, S2c and S2f, does not imply that the species will be able to expand its range in arbitrarily steep gradients. Natural selection imposes a phenotypic load on the mean growth rate of the population, given by the term 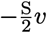 in (19), which increases proportionally with the inflation in trait variance as the environmental gradient becomes steeper. At exceedingly steep gradients, the phenotypic load becomes so strong that brings the population to extinction. In the absence of Allee effect, a maximum steepness of the gradient in trait optimum is given in [24] as 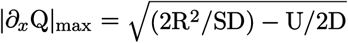, beyond which the population cannot survive—unless possibly marginally at the vicinity of the habitat boundary. This estimate of the critical gradient steepness for species’ survival is still approximately valid after our addition of the Allee effect to the model. Note that, the results in [24, Fig. 4] and [30, Fig. 2a] show that the range expansion speed also vanishes to zero exactly at this critical gradient, and remains nonzero at shallower gradients. This further implies that in linearly changing environments, genetic swamping cannot establish range limits when trait variance is free to evolve [25, 24].

Throughout the present work, by “maladaptive”, “disruptive”, or “swamping” effects of asymmetric core-to-edge gene flow we refer to the dispersal effects such as those shown in Figure 1d.

That is, we refer to the deviation of trait mean at peripheral populations due to receiving maladapted phenotypes from the core, often known as migration load. In the results presented in next sections we show how these maladaptive effects of gene flow are intensified by the interactions with interspecific competition, leading to formation of range limits, as in [20, 24]. We also show how these effects play the key role in stabilizing range limits at environmental knees and wiggles.

### Interspecific Competition, Asymmetric Gene Flow, and Formation of Species’ Range Limits

Interspecific competition and gene flow along an environmental gradient jointly contribute to the evolution of character displacement and range limits when two initially allopatric species become sympatric over a habitat region. For competitively identical species, this was first shown by Case and Taper [20] under the constant (in time and space) trait variance assumption, and then confirmed by Shirani and Miller [24] for the case where trait variance is free to evolve. Since understanding the basic underlying mechanism of range limits formation is important for better understanding the stabilization mechanisms we describe in next sections, here we briefly demonstrate the range evolution of two competitively identical (equal) species in a linear environment. We initialize the two species with the same parameters but at opposite sides of the habitat, and compute their range evolution under different levels of the steepness of the environmental gradient. The results are shown in Figure 2 and the supplementary Figure S4.

When the environmental gradient is sufficiently steep, we observe (Figure 2) that range limits are formed at the interface between the two species. The initially allopatric species adapt to new habitat locations and expand their range. They become sympatric over a small region when they first meet at the middle of the habitat. Since both species are relatively well-adapted to the environment when they meet, their individuals over the region of sympatry have similar phenotypes. This initiates strong interspecific competition between the individuals which creates character displacement within the region of sympatry. The character displacement further implies departure of the trait mean of each species from the trait optimum. The resulting maladaptation, along with the effects of interspecific competition, substantially decreases the population density of the two species at their interface (Figure 2a). This decline in density further increases the asymmetry in the core-to-edge gene flow within each species—from their populous center to their low-density peripheral populations at the interface of the two species. This asymmetric gene flow is maladaptive and, due to the term 2 D_*i*_*∂*_*x*_ (log *n*_*i*_) *∂*_*x*_*q*_*i*_ in the trait mean equation (6), moves the species’ trait mean further away from the optimum trait within the region of sympatry.

As the spatial overlap between the two species increases, the reinforcing feedback between character displacement and asymmetric gene flow continues to induce further maladaptation and a decline in density. Eventually, the level of maladapted phenotypes at each species’ interfacing edge with the other species becomes so strong that it prevents further local adaptation and range expansion. As a result, a range limit is established for each species and a region of sympatry is formed between the species at the middle of the habitat. The highlighted curves in Figures 2a and 2b show the steady-state population densities and trait means, respectively. Similar to the single-species simulation in Figure 1, the effects of dispersal and local effects of selection and competition on rate of change of trait mean are shown (for the first species) in Figures 2c–2e. Note that, in comparison with the single-species case (with the same gradient), the local adaptation rate by natural selection remains almost the same (Figure 2d vs. Figure 1e), whereas the maladaptive effects of asymmetric gene flow is substantially intensified (Figure 2c vs. Figure 1d). Additional details, including the spatial evolution of trait variances, can be found in a similar simulation shown in [24, Figure 7]. We further note that, although not shown here, intraspecific trait variance declines within each population over the region of sympatry. This reduced variation in phenotypes allows for strong competition, character displacement, and maladaptation; as required for the establishment of the range limits described above.

When the environmental gradient is not sufficiently steep, the maladaptive effects of asymmetric gene flow cannot reinforce character displacement to the level required to stop range expansion of the species within a finite habitat. As Figure S4 shows, in this case the overlap between the species continues to expand and the species eventually become completely sympatric over the entire available habitat. The evolution of this complete sympatry is, however, much slower than the evolution of the limited sympatry as in Figure 2a.

It is worth noting that the evolution of character displacement requires the diversifying (disruptive) selection generated by competitive interactions between phenotypes to be stronger than stabilizing selection [20]. The character displacement shown in Figures 2b and S4b becomes more pronounced as the strength of stabilizing selection decreases [20, Figure 5]. This is because greater character displacement reduces competition, providing a fitness advantage. On the other hand, greater character displacement implies greater deviation from trait optimum and hence larger fitness load imposed by selection. When selection is weak, the balance between these opposing forces on fitness is reached at greater extents of character displacement. A similar effect is observed if competition is intensified by increasing V_1_ and V_2_. The simulations performed in [24] for faster growing species (R_1_ = R_2_ = 2 T) also show a greater extent of character displacement. Here, we perform our simulations with (relatively) slowly growing species to follow our conceptualization of the range evolution of montane birds.

**Figure 5:**
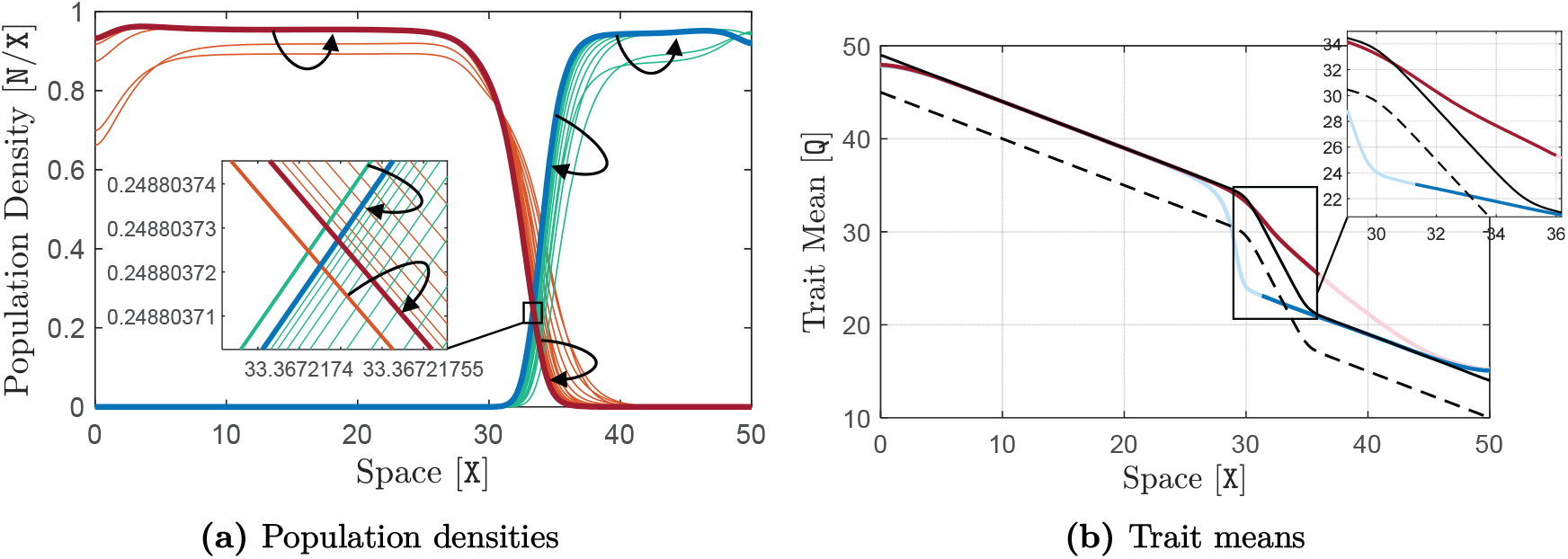
Robustness of the stability of the range limits against moderate climatic disturbances. The simulation is performed using the same model parameters as used in Figure 3. The equilibrium curves obtained at the end (*t* = 1500 `T`) of the simulation shown in Figure 3 are used as the initial curves here. Starting from *t* = 0 `T`, the curve of trait optimum Q is gradually shifted up by 4 `T` over a time course of 10 `T` and remains unchanged afterwards. That is, we add to Q a disturbance *δ*Q of the form shown in Figure (*iv*) in Box 2, with amplitude *α* = 4 Q and rise time *t*_r_ = 10 `T`. The initial curve of Q is shown by the dashed black line in (b), and the completely shifted curve after *t* = 10 `T` is shown by the solid black line. The simulation is performed for a time horizon of *T* = 500 `T`. Curves of population density are shown at every 6 `T` in (a), and the final curves obtained at *t* = 500 `T` are highlighted. The final curves of trait mean obtained at *t* = 500 `T` are shown in (b). The same description as given in Figure 2 holds for the curve colors. We observe that the range limits remain stable under the climate-warming disturbance applied at the beginning of the simulation. That is, the limits converge back to the same initial equilibrium after the time course of the perturbation is complete.

### Range Limits in Linear Environments: Stable or Unstable?

Our results show that in a linear environment (see Box 1), the range limits formed for competitively identical (equal) species in Figure 2a are evolutionarily stable in a weak sense (see Box 2). If, for example, we initialize the downslope species at the same location as in Figure 2a, but displace the initial population of the upslope species upward, then the distribution of the two species will converge to a different equilibrium, no matter how small the displacement in the initial population of the upslope species is. This new equilibrium will look essentially the same as the equilibrium shown in Figure 2a, but with a region of sympatry and range limits that are shifted upslope (see Figure (*vi*) in Box 2). Therefore, depending on where the populations are initially centered, their equilibrium range limits will settle at different locations. This further implies that the evolutionary range dynamics of the two species has a continuum of equilibria, shown schematically in Figure (*i*) in Box 2.

The weakly stable range limits formed in Figure 2 are not robust against environmental disturbances (Box 2) such as climate change that cause permanent changes in the trait optimum, no matter how small the disturbance amplitudes are. To see this, we initialize our simulation with the equilibrium populations of Figure 2a, and then gradually shift up the line of trait optimum by *α* = 2.5 `Q` over a time course of *t*_r_ = 10 `T`; see the schematics shown in Figure (*iv*) in Box 2. We then keep the trait optimum unchanged for the rest of the simulation (*T* = 500 `T`). We can conceptualize this environmental disturbance, for example, as a permanent change in the environment due to climate warming. The simulation results are shown in the supplementary Figure S5. We observe that, due to the climate-warming perturbation, the equilibrium distribution of Figure 2a is shifted upslope, resulting in a shift in the species’ range limits. Since *∂*_*x*_Q = −0.5 `Q`*/*`X`, the spatial shift in the range limits due to the 2.5 `Q` shift in the trait optimum appears to be exactly equal to 2.5*/*0.5 = 5 `Q`. This direct proportionality further implies that the weakly stable range limits formed in Figure 2a will not be restored to their original location when a permanent change occurs in climate, no matter how small the change is. It should be noted though, that the original range limits will be restored if the environmental disturbances appear as transient fluctuations (see Box 2), such that the trait optimum line is temporarily shifted up or down but eventually returns exactly to its original value. Such transient disturbances that completely vanish in time, unless large in amplitude, may not be of particular interest in studying stability of range limits.

#### Box 2

**Notions of stability**

We use standard notions of stability. Although the dynamics of the evolutionary PDE model we use is understood in an infinite-dimensional function space, we avoid the technical language of indefinite-dimensional dynamical systems theory [44, 45, 46]. The equivalent notions from finite-dimensional dynamical systems [47, 48, 49] are sufficient for conceptual understanding of our results.

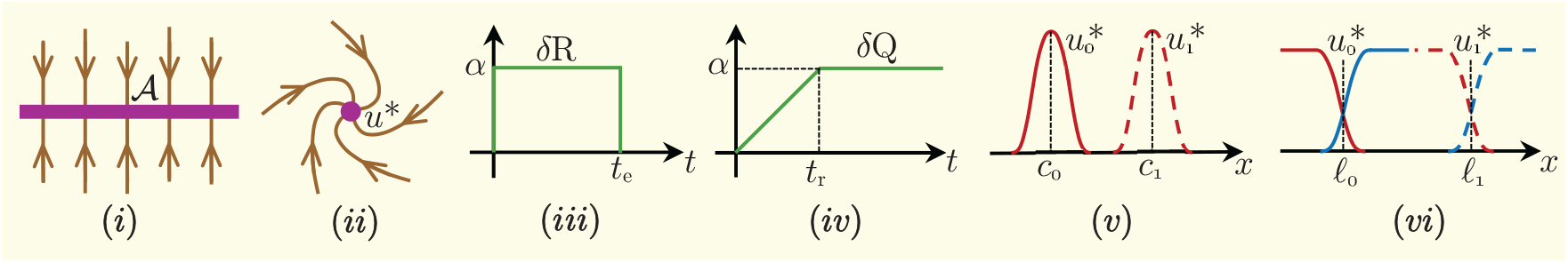

**Equilibrium**

An *equilibrium point*, or simply an *equilibrium* of a dynamical system is a steady state (fixed point) of the system: if the state of the system is initiated at an equilibrium, then it stays at the equilibrium for all time. For the PDE model that we use, a “point” is understood as a vector of functions (curves) in a space of all possible vectors of functions (e.g., curves of population density in Figures (*v*) and (*vi*)). An equilibrium is *isolated* if it has a neighborhood in which there does not exist another equilibrium. Otherwise, the equilibrium is *non-isolated*.

**Perturbations and disturbances**

An equilibrium state of a system is established for a fixed (constant) set of system parameters and external forcing terms. Yet the parameters are often subject to *perturbations*, and the system receives external *disturbances*. An *additive* perturbation *δ*R in a parameter R, for example, changes R to R + *δ*R. We say a perturbation is *transient* if it vanishes in finite time or asymptotically as *t* → ∞. Otherwise, we say the perturbation is *permanent*. The rectangular-shaped perturbation in Figure (*iii*), for example, is a transient perturbation with amplitude *α* and ending (termination) time *t*_e_. The disturbance *δ*Q in Figure (*iv*), with rise time *t*_r_ and amplitude *α*, is a permanent disturbance.

**Attracting set**

A set of points 𝒜 is *attracting* if there exists a neighborhood 𝒩 of 𝒜 such that all trajectories (evolution of system states in time) starting from the points in 𝒩 eventually converge to 𝒜 as *t* → ∞. The set 𝒜 (in purple) in Figure (*i*) gives an example.

**Stability**

An equilibrium is *(Lyapunov) stable* if all trajectories starting near the equilibrium stay close to it for all time. More precisely, an equilibrium *u*\* is stable if for any neighborhood 𝒰 of *u*\* there exits another neighborhood 𝒩 of *u*\* such that all trajectories starting in 𝒩 remain in 𝒰 for all time. Note that a stable equilibrium is not necessarily attracting, and it may or may not be isolated. An equilibrium that is both stable and attracting is *asymptotically stable* (Figure (*ii*)). An asymptotically stable equilibrium is isolated. When an equilibrium is stable but is not attracting, we say it is *neutrally stable*. A particular case of neutrally stable equilibria is where a system has an attracting set 𝒜 composed of a continuum set of (infinitely many) equilibria, such as the set 𝒜 shown in Figure (*i*). Note that although 𝒜 is an attracting set, none of the equilibrium points in 𝒜 is attracting. Every neighborhood of an equilibrium 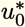 in 𝒜, no matter how small it is, contains another equilibrium 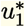. Trajectories starting form 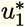 will not converge to 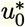. To which equilibrium trajectories starting from the *domain of attraction* (attraction neighborhood) of 𝒜 will converge depends on the initial state (starting point) of the trajectory. Note also that, when the system of Figure (*i*) is at an equilibrium, a small perturbation (force) to the system causes the state of the system to transition to another equilibrium.

**Weakly stable range limits**

We say a species’ range, and correspondingly its range limit(s), is *evolutionarily stable in a weak sense*, or *weakly stable*, if the species’ population dynamics (and hence its population density) is at a neutrally stable equilibrium. The limited range of a solitary species formed in a linear environment due to the swamping effects of gene flow, as identified in [26], is an example of a weakly stable range. Figure (*v*) shows, schematically, the equilibrium population densities *u*\* corresponding to such a limited range formation. Depending on where the population is initially centered, the equilibrium population distribution will be centered at *c*_0_, *c*_1_, or any other locations on the geographic axis. Therefore, there exists a continuum of equilibrium population states with limited range, similar to the case shown in Figure (*i*). Transient perturbations of small amplitude will shift the species’ equilibrium range from one location to another location. Similarly, the equilibrium population density curves shown schematically in Figure (*vi*), corresponding to the range limits formed between two competitively identical species, are weakly stable. The stable range limits shown by [20], or in Figure 2 of the present work, serve as examples.

**Strongly stable range limits**

If a species’ population dynamics is at an asymptotically stable equilibrium, we say the species’ range (or the range limits) are *evolutionary stable in a strong sense*, or *strongly stable*. Transient perturbations of sufficiently small amplitude, although temporarily move the species range limits, will not cause a permanent shift in the species’ equilibrium range. Once the perturbation vanishes, the species’ population will return back to the same equilibrium state it was initially at before the perturbation occurred. As we show, the stability of the range limits formed at environmental knees and wiggles is strong.

Although the weak stability of the range limits formed in Figure 2 makes them quite sensitive to disturbances, the crucial drawback of such range limits is that they are not generic. That is, in a (almost) linear environment, a coevolutionary equilibrium state such as the one shown in Figure 2a—with a fraction of each species’ range being sympatric and the remaining being allopatric— does not exist, almost surely. Our simulations show the existence of such equilibrium states only for the very specific case of “identical” (or “equal”) competitors, which is unlikely to ever be the case in the real world. When one of the species is competitively stronger than the other, it constantly pushes the interface between the two species towards the weaker species. This means that, an equilibrium state such as the one shown in Figure 2 will not form or, in other words, the competitively formed range limits will not be evolutionarily stable. In fact, since the environment is linear, every point in the habitat imposes the same level of selection pressure on both species. Therefore, the competitively weaker species does not receive any advantage over the other species from abiotic environmental factors; which could counterbalance the competitive difference between the two species and stabilize the range limits. To demonstrate the instability of the range limits for unequal competitors via simulations, we initialize the populations with the equilibrium curves of Figure 2 and increase each of the parameters R_1_, D_1_, V_1_, and K_1_ of the downslope species by only one percent. The simulation results, shown in the supplementary Figure S6, verify the instability of the range limits and their constant shift towards the weaker species.

In the real world, the difference between the competitive strength of two related species can be significant. As in [24], our simulations show two possible evolutionary outcomes of species range evolution in a linear environment: complete exclusion of the weaker species, or marginal existence of the weaker species at the vicinity of the habitat boundary. When one of the species is much stronger than the other, it completely excludes the other species. This is shown in supplementary Figure S7, where the downslope species is dominant due to its significantly larger maximum growth rate; R_1_ = 1.2 `T`^−1^ versus R_2_ = 1 `T`^−1^. When the difference between the two species is less significant, as in Figure S8, an evolutionarily stable state of marginal coexistence can eventually form as the interface between the two species approaches the habitat boundary. In this case, the weaker species survives with a limited density at close vicinity of the boundary. This is because when this species is eventually pushed against the boundary, it starts gaining advantage from its range contraction and density decline. Since there is no influx of phenotypes from the boundary, the contraction in range of the weaker species results in less migration load [43] to be created by gene flow in the peripheral population of this species (at its interface with the stronger species). Additionally, the decrease in density reduces the intraspecific competition load on the population. The stronger species, however, does not enjoy these advantages. As a result, the decrease in the fitness load on the weaker species compensates for this species’ moderate weakness in interspecific competition and allows the species to survive.

We conclude from our discussion above that, when the environment changes almost linearly in space, range limits formed by the interaction between interspecific competition and gene flow are unlikely to be evolutionarily stable. Below, we show how environmental knees and wiggles, as special types of plausible environmental nonlinearities, can strongly stabilize such range limits. We further show that the stability of the equilibrium range limits formed at these environmental nonlinearities is sufficiently robust against permanent environmental disturbances such as climate warming.

### Environmental Knees and Wiggles Can Stabilize Range Limits

The key nonlinearity in the environment that effectively stabilizes competitively formed range limits is a sharp change in the steepness of the gradient in trait optimum. In its simplest form, such a nonlinearity appears as a single knee in the environment. In its possibly more pervasive form, the nonlinearity may appear as two adjacent knees, forming a wiggle (see Box 1). We should note that changes in the spatial profile of the trait optimum at environmental knees and wiggles do not necessarily associate with a change in the productivity of the individuals. Therefore, the effects of knees and wiggles on species’ range dynamics described here are distinct from those of gradients in productivity.

We first show that environmental knees can stabilize the range limits formed between two species, which are otherwise unstable in a completely linear environment. For this, we simulate an environmental trait optimum that is linear everywhere, except at a knee located at *x* = 35 `X`. The steepness of the gradient changes sharply at the knee from 2.5 `Q`*/*`X` to 0.5 `Q`*/*`X`; see the black line in Figure 3b. We let the downslope species be stronger, with R_1_ = 1.1 `T`^−1^ versus R_2_ = 1 `T`^−1^. We initialize both species on the same side of the habitat, downslope of the knee, and let their range evolve for an evolutionary time horizon of *T* = 1500 `T`. The results are shown in the upper panel of Figure 3. We first note that in Figure 3a the species’ population density over the steep (downslope) part of the habitat is substantially lower than the environment’s carrying capacity. As we described before for a solitary population, this is because the phenotypic load imposed by natural selection is strong when the gradient is steep. However, the steepness 2.5 `Q`*/*`X` we chose for this part of habitat is still sufficiently smaller than the critical survival steepness |*∂*_*x*_Q |_max_ calculated in [24]. Based on the estimates provided in [24], an environmental gradient of 2.5 `Q`*/*`X` is biologically plausible as well, though it is very steep.

Figure 3a shows that the species’ competitively formed range limits converge to an equilibrium state near the environmental knee. Initially, the upslope species expands its range in both directions; both downslope towards the stronger species and upslope towards the knee. When this species crosses the knee, it experiences a much shallower gradient (0.5 `Q`/`X`) and hence much weaker migration and genetic loads. As a result, it grows to high density over this part of the habitat and expands its range relatively quickly up to the habitat boundary. On the other side, when this species meets the downslope species, a region of sympatry is initially created between the two species. Since the downslope species is stronger, it pushes the interface between the two species towards the knee. The interface is eventually stabilized at an equilibrium near the knee. The curves highlighted in red and blue in Figure 3a show the equilibrium (steady state) population distribution of the species.

The mechanism of range limits stabilization at an environmental knee relies on the contrast that the sharp change in the gradient’s steepness creates in the level of maladaptive gene flow to species’ peripheral populations. As the interface between the two species approaches the knee, the levels of asymmetric gene flow from central to peripheral populations of the two species change differently. The central population of the upslope species has already established itself in the region upslope the knee, and hence is well-adapted to the shallow gradient there. As a result, the peripheral population of this species suffers less disruptive effects (migration load) of gene flow as the interface approaches the knee. By contrast, the central population of the downslope species is adapted to the steep gradient downslope of the knee. Therefore, the phenotypes brought by gene flow to a local peripheral population of the downslope species are significantly greater than the average local phenotype. As a result, the trait mean of the downslope species significantly deviates (upward) from the trait optimum near the knee, causing a substantial fitness load on the peripheral population of this species. The curves of trait means in Figure 3b show the resulting difference between the (steady-state) local adaptation of the peripheral populations of the two species. Since competition between the two species takes place within their overlapping peripheral populations, the advantage that the upslope species gains from reduced maladaptive effects of gene flow eventually compensates for its competitive weakness. This balances the competition between the two species and stabilizes their range limits as an equilibrium near the knee.

The coevolutionary mechanism of range stabilization we described above suggests that the competitively formed range limits at environmental knees are evolutionarily stable in a strong sense (see Box 2). The peripheral populations of the two species at their steady-state interface formed near a knee are subject to two opposing forces. If a transient perturbation slightly moves the interface upslope, then the fitness advantage that the upslope species has due to its local adaptation to the environment upslope of the knee gives this species the competitive strength needed to push the peripheral population of the downslope species (and hence the interface) back to the equilibrium location. If the interface is moved downslope by a perturbation, then the intrinsic competitive dominance of the downslope species works to push the interface back to the equilibrium location. Therefore, the presence of these two opposing forces ensures that the equilibrium state (point) of the species’ range evolution is isolated (see Box 2). Unlike the weakly stable range limits in Figure 2, the range dynamics shown in Figure 3a also implies that the equilibrium population distributions are not very sensitive to where the populations are initially centered. We provide additional simulations below, for the similar case of range stabilization at environmental wiggles, that further demonstrate the strong stability of the range limits formed at environmental knees.

To demonstrate the range stabilization dynamics at environmental wiggles, we simulate an environmental trait optimum that is linear everywhere with slope −0.5 `Q` */* `X`, except at a wiggle where its slope switches sharply to −2.5 `Q` */* `X`; see the black line in Figure 3d. Same as the simulations we performed above for an environmental knee, we let the downslope species be stronger, with R_1_ = 1.1 `T` ^−1^ versus R_2_ = 1 `T` ^−1^. We initialize both species on the same side of the habitat, downslope of the wiggle, and compute their range evolution for *T* = 1500 `T`. The results are shown in the lower panel of Figure 3. We first note that despite the sharp transition in the steepness of the environmental gradient at the wiggle, the upslope species can still adapt to and cross the wiggle; thereby expanding its range indefinitely up to the upslope habitat boundary. This means that the wiggle is not a physical barrier to range expansion.

Figure 3c shows that the spatial distribution of the two species converges to an equilibrium, at which the species’ range limits is stabilized near the upslope knee of the wiggle. When the interface formed between the two species enters the wiggle’s transition zone, the substantially steeper environmental gradient of the wiggle intensifies the interactions between species’ population density and gene flow. These intensified interactions enhance character displacement within the wiggle, sharpen the interface between the species, and slow down the advancement of the interface. The interface is eventually stabilized near the upslope knee of the wiggle.

The mechanism through which environmental wiggles stabilize range limits is essentially the same as the stabilization mechanism at environmental knees. In fact, only one of the knees of the wiggle (the upslope knee in Figure 3) plays the main role in stabilizing the limits. The steepness change at the other knee provides a further support for the stability of the limits, depending on how close the two knees are to each other. In the results shown in Figure 3c, the downslope species can grow to high density in the moderate gradient below the downslope knee. Since the wiggle is relatively short, the disruptive effects of asymmetric gene flow from the high-density core of this species still remain in effect in its peripheral population. The increased migration load on this population, which is in competition with the other (intrinsically weaker) species near the upslope knee, then increases the chance of species’ range stabilization. The effects of the downslope knee of the wiggle on increasing the migration load on the peripheral population of the downslope species can also be verified by comparing the steady-state levels of character displacement (difference between trait mean curves) in Figures 3b and 3d. The extent of character displacement at *x* = 35 `X` in Figure 3b (where the single environmental knee is located) is equal to 4.83 `Q`, whereas as the character displacement at the same location in Figure 3d (where the upslope knee of the wiggle is located) is equal to 5.24 `Q`.

Since range stabilization mechanisms at environmental knees and wiggles are essentially the same, our previous discussion on the stability of the range limits at environmental knees also suggests that the range limits formed at wiggles are evolutionarily stable in a strong sense (Box 2). Due to the opposing eco-evolutionary forces acting on the interface of the two species when it gets stabilized at the wiggle, the equilibrium state that the species’ range evolution converges to is isolated. Convergence to this equilibrium is fairly insensitive to the initial distribution of the two species. Supplementary Figure S9 shows, for example, that the population distributions converge to the same equilibrium as in the lower panel of Figure 3 even in the case where the two species are initialized at opposite sides of the wiggle. Moreover, simulations of transient perturbations (see Box 2) in each of the characteristic parameters of the species confirm that the equilibrium state of Figure 3 is precisely restored after being temporarily disturbed by the perturbations. See, for instance, the results shown in Figure S10 for additive perturbations of 20 percent in amplitude that last for 200 T in each of the parameters R_1_, D_1_, V_1_, and K_1_ of the downslope species. In the results presented in the next section we additionally show that the stability of the strongly stable range limits formed at wiggles (as well as knees) is fairly robust against permanent climatic disturbances.

We conclude this section by exploring, to some extent, the parameter values that make a wiggle effective enough to stabilize the range limits. In general, whether or not a wiggle will stabilize the interface formed between two competing species depends on all characteristic parameters of the species that affect their competitive strength relative to each other, the steepness and width of the wiggles, and the steepness of the environmental gradient outside the wiggle. These essentially include all parameters of the model and the wiggle we have used in our simulations. Since the comprehensive exploration of the entire multi-dimensional parameter space of the model is infeasible, we only focus on the effects of the steepness and width of the wiggles. Specifically, we set the parameters of the species equal to those used in Figure 3, and fix the slope of the environment outside the wiggle at −0.5 `Q`/`X`. We then let the width and steepness of the wiggle change over a reasonably wide range of values, and identify the critical values of width and steepness at which a transition (bifurcation) occurs in the wiggle’s effectiveness. We perform this analysis for two different competition strengths, determined by the values of phenotype utilization variances. The results are shown in Figure 4.

For the given parameter values, the wiggles shallower or narrower than the ones on the curves shown in Figure 4 cannot stabilize the range limits, whereas the wiggles steeper or wider than those can. When competition between the species is more intensive (larger values of V_*i*_), the range limits can be stabilized by a shallower or a narrower wiggle. The results shown in Figure 4 suggest that narrow wiggles, with a width approximately less than 2 `X`, can be sufficiently effective in range stabilization only if they are very steep. This is because the width of narrow wiggles is comparable to the mean dispersal distance of the individuals in one generation time. As a result, a significantly large portion of the peripheral population of the stronger species (downslope species here) will cross a narrow wiggle by dispersal, before being eliminated by natural selection. Unless this population receives a substantial level of maladaptive gene flow due to the wiggle being very steep, it will be able to outcompete the weaker species in the shallow environment adjacent to the wiggle and push the range limits out of the wiggle. Sufficiently wide wiggles, on the other hand, can stabilize range limits even if they are only moderately steep; for example, with a steepness as low as 0.9 `Q`/`X`, compared with the outside-wiggle steepness of 0.5 `Q`/`X` used in the results of Figure 4. We see further that, when the wiggles are sufficiently wide, their minimum steepness required for range stabilization is almost independent of their width. This is because in wide wiggles only one of the knees (upslope knee here) plays the essential role in stabilization. However, as we discussed for the results shown in Figure 3, close adjacency of the knees can result in a more pronounced character displacement at the stabilizing knee of the wiggle. Greater character displacement then provides greater robustness for the stability of the range limits against environmental changes, as we discuss below. Based on these observations, the curves shown in Figure 4 suggest that a wiggle width approximately equal to 4 `X` is almost optimal for the species’ competitive difference that we simulated: it is wide enough to allow for lower stabilizing steepness, and is narrow enough to provide better robustness against disturbances. The wiggle width 5 `X` that we used in our other simulations is close to this optimal value.

### Climate Change and Stability of the Range Limits

The strong stability of the range limits formed as a coevolutionary equilibrium at environmental knees and wiggles is robust against permanent climatic disturbances, provided they are sufficiently small in amplitude. Here, we only show the robustness at wiggles, noting that the robustness at knees hold similarly. Moreover, we only consider permanent climate-warming disturbances; robustness against climate-cooling disturbances are similar, but with the “upslope” and “downslope” attributes interchanged. We present the results for a decreasing environmental optimum, that is, the trait optimum decreases as species move upslope. If individuals are represented by a trait that increases with elevation, such as wing length in birds [50], the same results hold but with interchanging the “upslope” and “downslope” attributes.

To perform our analysis, we initialize our simulation with the two equilibrium populations of Figure 3, whose range limits have been stabilized at the wiggle. We simulate a climate-warming disturbance by gradually shifting up the curve of trait optimum Q by an amplitude *α* = 4 Q over a time course (rise time) of *t*_r_ = 10 T, starting from the beginning of the simulation. When the course of climate warming is complete, we fix the curve of Q at its new profile until the end of the simulation. See Figure (*iv*) in Box 2 for an schematic of this environmental disturbance, *δ*Q. We allow the species to evolve for *T* = 500 `T`. The results are shown in Figure 5.

The range dynamics shown in Figure 5 confirms that the stability of the range limits is robust against the climate-warming disturbance that we simulated here. At the beginning of the simulation, both species suffer some population losses as they fail to fully adapt to the quickly changing environment. The peripheral population of the downslope (stronger) species initially has trait mean values greater than the environmental optimum, particularly near the upslope knee of the wiggle where the range limits get stabilized. Therefore, this population suffers less from the incremental changes in the optimum phenotype, and in fact enjoys the changes at some locations. As a result, the range limits of the two species are slightly pushed upslope by this species when climate warming is taking place. However, once the warming is complete and both species successfully adapt to the new environment, the range limits return to their initial equilibrium state.

Climate warming can still destabilize the stable range limits in Figure 3 if it is sufficiently large in amplitude and occurs sufficiently fast. To see this, we repeat the same simulation as described above (Figure 5), but with a stronger climate-warming disturbance of amplitude 8 Q. The evolution of the species’ range for 100 `T` is shown in Figure 6a. Here, the advantage that the peripheral population of the downslope (stronger) species has—due to its partial pre-adaptation to larger phenotype values within the wiggle—is significant enough to allow this species push the range limits entirely out of the wiggle, before the upslope species finds a chance to sufficiently adapt to the warming climate. As a result, the range limits become destabilized and the downslope species succeeds in crossing the wiggle and expanding its range.

**Figure 6:**
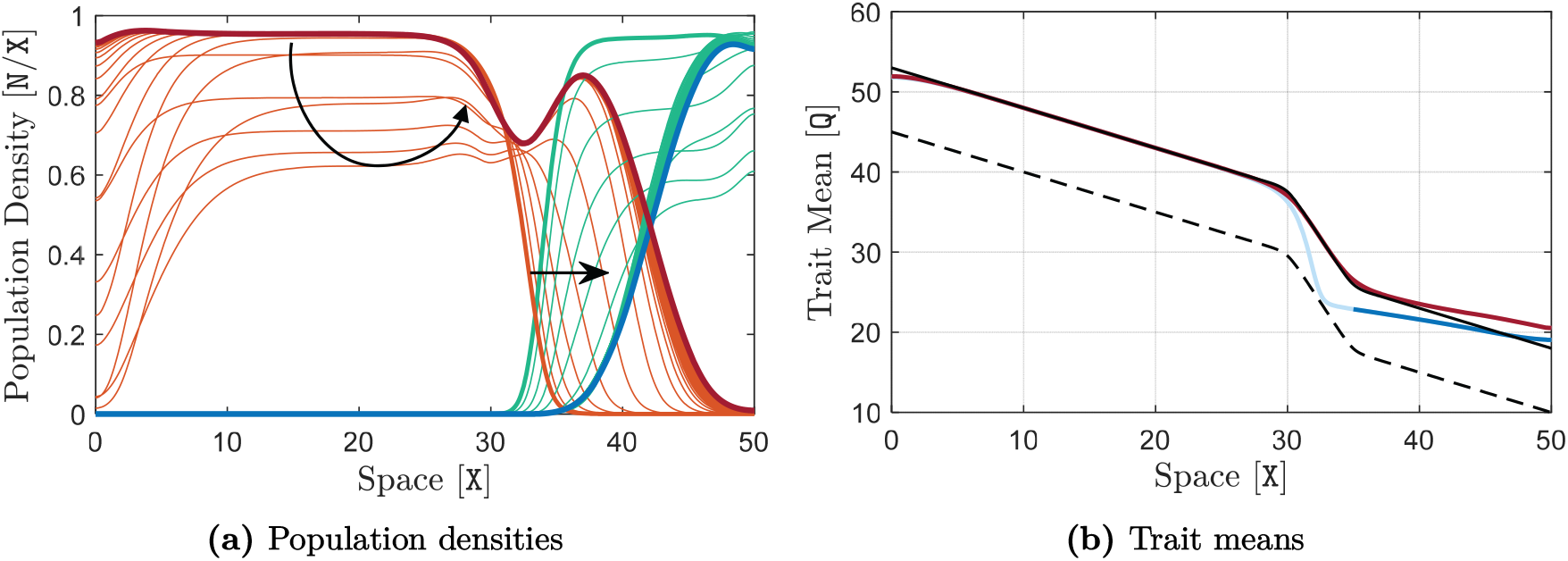
Destabilization of the range limits by a strong climate-warming disturbance when the downslope species is stronger. The same simulation as presented in Figure 5 is performed here, with the only difference being that the shift in the trait optimum takes a larger amplitude of *α* = 8 `Q`. The curves of population density are shown in (a) at every 2 `T`, up to the time *t* = 100 `T`. The final curves are highlighted in red and blue at *t* = 100 `T`, and their corresponding trait mean curves are shown in (b). The same description as given in Figure 5 holds for curve colors and the dashed curve. Arrows indicate the direction of evolution in time. Due to the larger amplitude of the climate-warming disturbance, the initially stable range limits are destabilized, resulting in an upslope expansion in the stronger species’ range. Further continuation of the simulation up to *T* = 2000 `T` is shown in supplementary Figure S11, which shows that the weaker species will eventually be completely excluded from the habitat.

**Figure 7:**
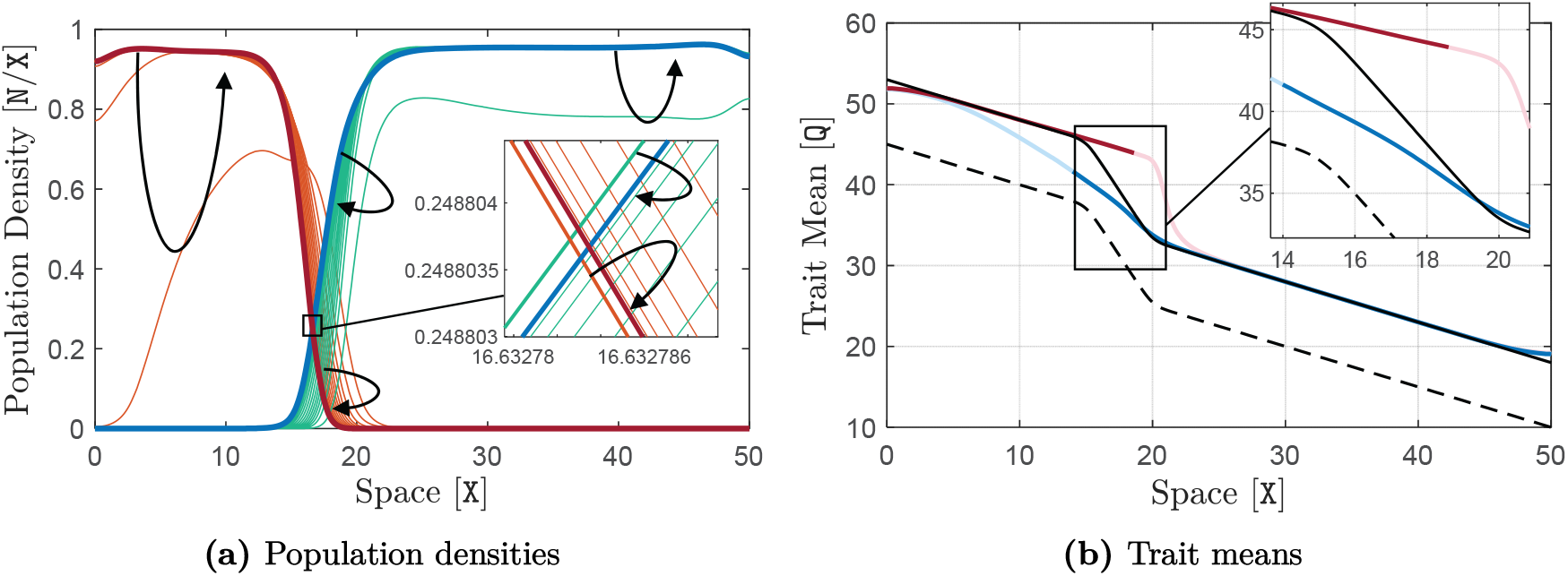
Robustness of the stability of the range limits against strong climate-warming disturbances when the upslope species is stronger. Here, R_1_ = 1 `T`^−1^ and R_2_ = 1.1 `T`^−1^, that is, in contrast with the results shown in Figure 3, the upslope (green) species is made stronger here. The rest of the parameters take their typical values given in Table 1. The environmental wiggle is located over the interval [15 `X`, 20 `X`]. The formation of the stable range limits near the downslope knee of the wiggle is shown in supplementary Figure S12. Similar to the simulation associated with Figure 6, the equilibrium curves obtained at the end of the simulation shown in Figure S12 are used as the initial curves here. The climate-warming disturbance *δ*Q is also applied in the same way as it was applied in Figure 6, with the same amplitude of *α* = 8 `Q`and rise time *t*_r_ = 10 `T`. The simulation is performed for a time horizon of *T* = 500 `T` and curves of population density are shown in (a) at every 12 `T`. The final curves obtained at *t* = 500 `T` are highlighted in red and blue, and their corresponding trait mean curves are shown in (b). The same description as given in Figure 6 holds for the curve colors, arrows and the dashed curve. We observe that, unlike the case where the downslope species was stronger (Figure 6), here the range limits remain stable even though the climate-warming disturbance is large in amplitude.

In general, there are several possibilities for the future range evolution of the species (with the downslope species being stronger) when their range limits formed at a wiggle are destabilized by climate warming. If there is another wiggle upslope to the previous wiggle, then the range limits can be stabilized at the new wiggle. In this case, the strong climate-warming disturbance indeed causes an upslope shift in species’ range limits from one wiggle to another wiggle. In the absence of another wiggle, the stronger species may eventually exclude the weaker one entirely from the habitat, or the species’ distribution may converge to an evolutionarily stable state of marginal coexistence at the vicinity of the habitat boundary; such as the state shown in supplementary

Figure S8. Our further continuation of the simulation of Figure 6 up to *T* = 2000 `T` shows complete exclusion of the weaker species; see supplementary Figure S11. Another possibility, which is not shown here, is that the species’ ranges converge to an evolutionarily stable equilibrium at which the stronger species occupies the entire available habitat but the weaker species still survives at the vicinity of the wiggle with a low density and limited range.

Finally, we show that if the upslope species is stronger, then the stability of the range limits formed at a wiggle is robust even against strong climate-warming disturbances. For this, we repeat the simulations of Figures 3 and 6, but this time with R_2_ = 1.1 `T`^−1^ versus R_1_ = 1 `T`^−1^. That is, we first initialize the two species upslope of a wiggle and let their ranges be stabilized at the wiggle. The results are shown in supplementary Figure S12. Then, we use the equilibrium populations of this simulation to initialize the second simulation, where we apply a strong climate warming disturbance *δ*Q of amplitude *α* = 8 `Q` and rise time *t*_r_ = 10 `T`, the same as what we did in Figure 6. We let the species’ range evolve for *T* = 500 `T`. The results are shown in Figure 7. Similar to what we saw in Figure 6a, during the course of climate warming the range limits are (slightly) pushed upslope by the downslope species whose peripheral population is partially pre-adapted to larger phenotypes. However, since this species is competitively weaker here, it cannot push the range limits out of the wiggle. Even if it can, possibly in the case of a stronger warming disturbance, the stronger upslope species will eventually push the limits back again to their original equilibrium in the wiggle (once the course of climate warming is complete and both species become adapted to the new climate).

We conclude our results by exploring the disturbance parameter values *α* and *t*_r_ that make a climate-warming disturbance strong enough to destabilize the range limits formed at a wiggle; for the case where the downslope species is stronger and trait optimum is decreasing. For this, we repeat the simulation used in Figure 5 for different values of disturbance amplitude (*α*) and rise time (*t*_r_), chosen within a reasonably wide range of values. By checking the divergence (destabilization) of the range limits at the end of each simulation, we identify the critical values of these parameters: a disturbance that has higher amplitude or occurs faster than these critical values can destabilize the range limits. We perform our analysis for wiggles with three different levels of steepness but with the same width set equal to the almost optimal value 5 `X`. The results are shown in Figure 8.

**Figure 8:**
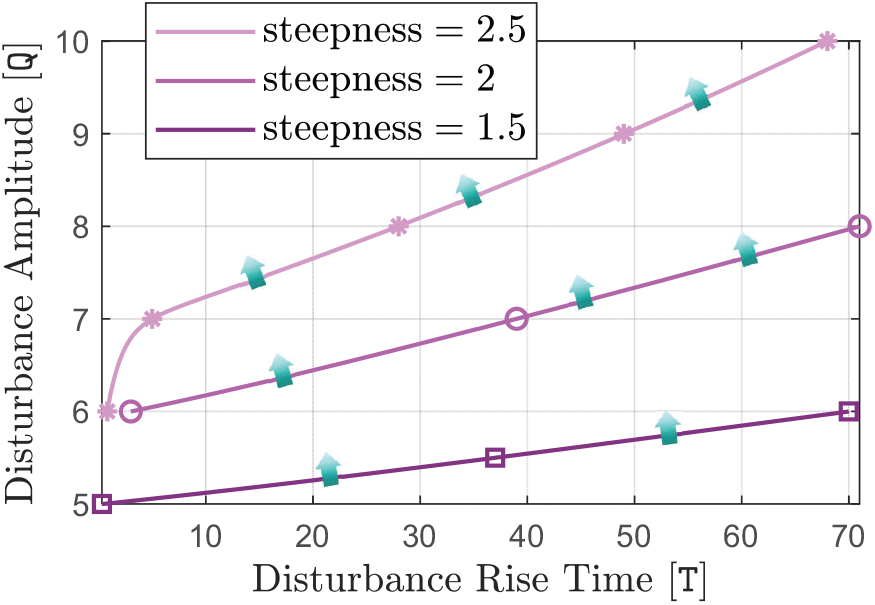
Critical climate-warming disturbances that can destabilize the range limits formed at environmental wiggles. The simulation associated with Figure 5 is repeated here for different values of the amplitude and rise time of the climate-warming disturbance, to identify the critical parameters that make the disturbance strong enough to destabilize the range limits formed at the wiggle. To ensure that the stability or instability of the equilibrium near the critical values is not affected by the habitat boundary condition, the habitat’s upslope boundary is extended to *x* = 70 `X`, with the wiggle’s location remaining unchanged. To find the critical disturbance parameters, approximately, different values are chosen for the disturbance amplitude within a reasonably wide range of values. A step-wise search (step-wise changes of step size 1 `T` in the rise time) is then performed at each amplitude value to find the critical value of the disturbance rise time, below which the disturbance can destabilize the range limits. To do the searches, the simulation is run at each step (i.e., for each (amplitude, rise time) pair) for a period of *T* = 500 `T` and divergence of the range limits from the initial equilibrium is checked at the end of the simulation. The critical parameter values obtained through the searches are marked in the graph. The critical values marked by asterisks are associated with simulations with the steep wiggle (steepness = 2.5 `Q`/`X`) used throughout the paper. The critical values marked by circles and squares are associated with a moderately steep and a shallow wiggle, respectively. The curves connecting the points are computed by interpolation. Disturbances that have higher amplitude or occur faster (have shorter rise time) than the critical values marked in the graph (lie above the curves) can destabilize the range limits. Arrows indicate transition from stable (dark) to unstable (light) range limits as the disturbance parameters change across the curves.

The curves of critical disturbance strength parameters in Figure 8 imply that steeper wiggles stabilize the range limits more strongly. That is, the stability of the range limits is more robust against permanent climatic disturbances when the wiggle is steeper. The steeper the wiggle is the stronger (more rapidly changing with higher amplitude) the disturbance must be to be able to destabilize the range limits. We further see, especially at shallow wiggles, that the destabilizing strength of the disturbance is more effectively determined by (is more sensitive to) its amplitude rather than its speed (rise time). In particular, no matter how fast the disturbance takes place, a minimum disturbance amplitude of about 5 `Q` is necessary to destabilize the range limits; a value comparable to the maximum level of character displacement 5.24 `Q` that we computed for the coevolutionary equilibrium state of the populations in the lower panel of Figure 3.

## Discussion

In this work we used a fairly comprehensive deterministic model of species range evolution [24] to analyze the stability of range limits formed by interspecific competition. We proposed and extensively studied environmental knees and wiggles (Box 1) as prevalent environmental nonlinearities that can robustly stabilize competitively formed range limits which are otherwise unstable in (almost) linear environments. We specifically identified the contrast created in the level of maladaptive gene flow to species’ peripheral populations—when their competitively formed interface reaches an environmental knee—as the key factor in stabilizing the range limits. This contrast originates from the sharp change in the steepness of the environmental gradient occurring at a knee.

We note that Case and Taper [20] also discussed that the interface between two competing species will often become “attached” to places where environmental nonlinearities in the form of wiggles (or “kinks” as Case and Taper call them) exist. However, the mechanism they describe for attachment of the range limits to wiggles is only valid for the non-generic and ecologically unlikely case where competitors are identical. The mechanism relies on the fact that the steep environmental gradient in a wiggle slows down the range expansion of the two species—initially located at opposite sides of the wiggle—thereby increasing the chance that they will meet within the wiggle. The range limits formed in this special case will be weakly stable (see Box 2). As also pointed out in [20], the equilibrium ranges formed at the wiggle through this “slowing-down” mechanism are sensitive to initial conditions. For example, if the two species are initialized at the same side of the wiggle as in our simulations, then the interface of the two (identical) species will simply stay at the location where the species first meet and will not move towards the wiggle. In contrast, the stabilization mechanism we identified in this work holds for the generic case of unequal species, results in range limits that are evolutionarily stable in a strong sense (see Box 2), and it essentially requires just a single knee in the environment (not necessarily a wiggle). This mechanism relies on the sharp change in the steepness of the gradient at a knee, rather than reduction in the rang expansion speed of the species over the transition (steep) zone of a wiggle.

### Implications for Range Limits and Their Stability

The results we obtained have implications for the ongoing debate in ecology about how and when interspecific competition limits species’ ranges [51, 52]. Our results confirm that the interaction between interspecific competition and gene flow along an environmental gradient can set range limits [20, 21, 24]. However, we argue that when the effects of environmental conditions and biophysical constraints on the species (particularly those affecting a trait optimum) vary almost linearly in space, the range limits are generically unstable: in the absence of physical barriers or additional interacting species, the interface between the species constantly moves towards the weaker species. This may eventually lead to complete exclusion of the weaker species, or to an evolutionarily stable state at which the weaker species marginally coexists with the dominant species at the vicinity of the habitat boundary; see supplementary Figures S7 and S8.

If the environment has a sufficiently steep wiggle ahead of the moving interface, or a knee at which the steepness of the environmental gradient changes significantly, then the moving limits eventually get stabilized at the wiggle or the knee. Environmental wiggles are likely more prevalent stabilizers of range limits, because species are less likely to colonize very steep gradients over a very wide habitat region near an isolated knee. Moreover, unlike solitary knees, wiggles serve as bidirectional range stabilizers: species interfaces moving from downslope areas to a wiggle will get stabilized near the upslope knee of the wiggle, whereas the interfaces approaching a wiggle from upslope areas will be stabilized near the downslope knee of the wiggle. If a wiggle is very narrow, of a width comparable to the mean dispersal distance of the species in one generation time, then it must be very steep to be able to stabilize the range limits; see Figure 4. The steepness of the wiggle must also be higher if the competitive difference between the species is greater. When the two species have more generalist individuals (larger values of V_*i*_), interspecific competition is intensified and range limits can be stabilized by a shallower or a narrower wiggle; see Figure 4.

We showed that the stability of the range limits formed at wiggles and knees is fairly robust against sufficiently small perturbations in the trait optimum, such as those caused by permanent climate change disturbances. The steeper the wiggle (or knee), the more robust the stability of the range limits; see Figure 8. Whether or not the range limits are destabilized by strong climate changes depends on the type of the change (warming or cooling) as well as the relative position of the species across the wiggle. In our conceptualization of a trait that is positively correlated with an elevation-dependent temperature gradient (that is, smaller (cooler) trait values at higher elevations) sufficiently strong climate-warming disturbances destabilize the range limits when the downslope species is dominant. In this case, the strong disturbance allows the dominant species to cross the wiggle and make an upslope expansion in its range. By contrast, the stability of the range limits is robust, even against strong climate-warming disturbances, when the upslope species is dominant. The converse direction of effects hold for climate-cooling disturbances, as well as the case where the trait is negatively correlated with temperature (increases with elevation).

We should note that whether a wiggle with given width and steepness will stabilize the range limits depends on the level of competitive difference between the species, as well as the steepness of the shallow gradient outside the wiggle. The competitive strength of the species relative to each other is determined by the combination of the effects of differences between species on each and any of their characteristic parameters. Such differences lie in a high-dimensional parameter space that cannot be quantitatively characterized. Therefore, our general observations discussed above (as in Figures 4 and 8) should be interpreted quantitatively rather than quantitatively.

### Random Dispersal and Maladaptive Core-to-Edge Gene Flow

The mechanism of range limits stabilization at environmental knees and wiggles we described in the present work relies on the maladaptive effects of asymmetric core-to-edge gene flow on adaptation of the peripheral populations. By maladaptive (swamping) effects of gene flow we mean the load that migration imposes on peripheral populations by bringing phenotypes that are adapted to the abundant core but maladapted at the edge, hence deviating the peripheral populations’ trait mean from the trait optimum; see Figures 1d and 2c. There is strong evidence that many species show declines in abundance towards their range edge; see for example [53, 54] for North American birds. Evidence for asymmetric core-to-edge gene flow has also been found in nature [55, 56, 57, 58, 59]. The findings in [55], in particular, are consistent with the center (core)-periphery hypothesis [60], that genetic variation and demographic performance of a species decrease from the center (core) to the edge of its range. The spatial patterns of trait variance and trait optimum in our results (Figure 1), as well as those shown in [24] are also consistent with this hypothesis.

Despite the available evidence, as discussed above, whether asymmetric gene flow consistently occurs from central to peripheral populations, and whether central-to-edge gene flow reduces mean fitness in edge populations, has been generally questioned [61, 62, 63, 60]. This is in part because patterns of gene flow depend on multiple factors, such as population size, environmental context, genetic variation, and dispersal behavior [62]. The methodologies used to uncover the patterns can also affect the outcomes.

One potential source of difficulty, and possible misinterpretation, in examining the effects of gene flow can be due to the contrasting effects of random gene flow: the swamping effects on population trait mean (Figure 1d) versus the enhancing effects on population adaptive potential (Figure 1e) [59, 62, 64, 61]. Our results, as well as those in [24] and [30], show that the range expansion speed of a solitary species decreases as the steepness of the environmental gradient (hence the level of gene flow) increases. This implies that, in the model we used, the overall effects of random gene flow on range expansion capacity are not facilitative. Disentangling the contrasting effects of gene flow on range expansion speed is not straightforward. Yet, we propose that the reduction in expansion speed could be, in part, due to the fact that trait variance evolves more slowly than the trait mean. Thus, the increased adaptive potential in steeper environments (due to the inflation in trait variance) cannot completely compensate for the swamping effects. We note that, interspecific competition intensifies the swamping effects of gene flow without significantly affecting the adaptive potential, resulting in the formation of range limits; see Figure 2.

Other reasons for the scarcity of evidence for significant maladaptive gene flow include that environmental gradients are often shallow, species exhibit reduced dispersal, or species disperse adaptively (non-randomly). When an environmental gradient is steep and dispersal is random, reduced dispersal can allow for faster range expansion by reducing maladaptive gene flow [65, 30]. There is also empirical evidence confirming that many species in nature disperse adaptively to better habitat [66, 67, 68, 69, 70]. A phenotype-dependent dispersal strategy such as matching habitat choice [71, 72] can indeed make gene flow adaptive [30, 73, 74, 75, 76]. However, such an adaptive dispersal strategy may evolve only when the environmental gradient is very steep [30].

When an environment has an isolated knee with a sufficiently wide region of steep gradient, the stable range limits formed at the knee may be destabilized if the stronger species evolves an adaptive dispersal strategy. Such a strategy releases the species from the competitive disadvantage of strong maladaptive gene flow to its interface with the other species. However, when range limits are stabilized at an environmental wiggle, evolution of dispersal strategies is less likely. Only the low-density peripheral population of the stronger species will occupy the steep gradient inside the wiggle, whereas the core of the species will reside in the shallow gradient outside the wiggle. Since the peripheral population constantly receives phenotypes from its core, which are adapted to the shallow gradient, it may not be able to evolve an effective dispersal strategy. This is a further reason why environmental wiggles might be more prevalent stabilizers of range limits, rather than isolated knees. In addition, it suggests that environmental wiggles could be the habitat locations where empirical studies aiming for finding maladaptive core-to-edge gene flow could focus on.

### Environmental Nonlinearities and Range Limits in Nature

The importance of interspecific competition in limiting species’ ranges in nature is ultimately an empirical question, but our theoretical results shed light on this question. In particular, we highlight the crucial role of sharp changes (knees) or wiggles along environmental gradients in determining where interspecific competition can set stable range limits. Nonlinearities along environmental gradients, such as wiggles, would seem to be common in empirical systems. For example, ecotones between habitats are commonly observed. Some empirical observations also support an important role for step-like changes in environmental gradients to stabilizing species’ range limits. For example, closely related species often share a common range border at ecotones [77, 78] and other transitional locations along environmental gradients [79]; consistent with the predictions of our analyses.

Our investigation of the environmental linearity and nonlinearity based on the optimum value that the environmental conditions impose on a trait allows for broader implications. The determination of the optimum trait is also subject to functional trade-offs and constraints [80, 81]. For example, biomechanical, biochemical or physiological trade-offs may affect the trait optimum during the stages of development. Such trade-offs may vary nonlinearly even though the abiotic environmental conditions (such as temperature) vary linearly in space. In particular, the nonlinearity in trade-offs may create a knee or wiggle in the trait optimum, hence the selective gradient it imposes on the population. Within the framework of our study, an environment with such nonlinearities in the trade-offs is considered nonlinear, even if the environmental conditions appear to be linear. Since the knees and wiggles caused by trade-offs impose the same nonlinearities in the selective gradient as we investigated throughout our work, the stabilization mechanism we described also hold for such nonlinearities.

The nonlinearities caused by functional trade-offs may respond differently to climatic changes in the environment. Assuming a temperature-correlated trait, in our simulations we modeled a climate-warming disturbance by shifting the whole profile of the trait optimum upward. The location of the wiggles remain unchanged under such a disturbance. When wiggles in the trait optimum are due to nonlinearities in trade-offs, a climate-warming disturbance could also move the location of the wiggle. If an environmental disturbance moves the wiggle towards the weaker species, then the range limits will shift and get stabilized at the new location of the wiggle, no matter how large the disturbance is; see supplementary Figure S13. If the disturbance causes the wiggle to move towards the stronger species, then the range limits will still get stabilized at the new wiggle location, provided the disturbance is sufficiently small; see supplementary Figure S14. However, if the disturbance is so strong that the wiggle moves rapidly and entirely inside the core of the stronger species’ population, then the range limits will get destabilized and the stronger species will push the weaker species to the habitat boundary. This is because in this case the disturbance effectively passes the stronger species across the wiggle.

### Species’ Response to Climate Change

Our results provide a perspective on how climate change will lead to changes in species’ ranges [82]. Most species are currently shifting poleward and upslope toward cooler environments, but many species have stable ranges or are even shifting towards warmer conditions [83, 84]. One possible explanation for these divergent responses to climate change is that species’ interactions shape their ranges along environmental gradients. For example, interspecific competition between closely related species of tropical birds is an important factor limiting their elevational ranges [17, 85, 86]. The simple expectation is that species’ ranges will shift upslope as environments get warmer. However, our results suggest that this would hold only if the downslope (lower-elevation) species is the dominant competitor. Empirical evidence demonstrates this is not always the case [87]. Therefore, our results emphasize the importance of considering the competitive dominance between species when predicting changes in their range limits. When the environmental trait optimum varies almost linearly, a stronger competitor can expand its range regardless of whether it is located downslope to the other species (leading to an upslope shift in ranges) or upslope to it (leading to a downslope shift in ranges). If the range limits of the species are already stabilized at an environmental wiggle, and their trait is positively correlated with temperature, then climate warming (if strong enough) can only result in an upslope shift in ranges, and that occurs only when the downslope species is dominant. If the species’ trait is negatively correlated with temperature, then climate warming can only result in a downslope shift in ranges, and that occurs only when the upslope species is dominant. These findings particularly support the importance of measuring competitive ability of species empirically [88, 53, 89] in order to predict species’ responses to climate change.

### Future Research Directions

The model we used in our study does not include the effects of stochastic evolutionary processes such as genetic drift, which are proposed to play an important role in setting range limits under certain conditions [90, 64]. The mean fitness of small and possibly isolated populations at range margins can be substantially impaired by the load of random genetic drift, that is, the reduction in population-mean fitness due to the fixation of deleterious mutations by genetic drift [28, 27]. Genetic drift can also erode genetic variation in small populations, resulting in a reduction in their adaptive potential. When competitively formed range limits are stabilized at an environmental wiggle (or a knee), as we described, the peripheral population of the stronger species may suffer a strong reduction in density along the steep gradient within the wiggle; see Figure 3. This makes the stronger species more sensitive to genetic drift at the environmental knee where it interfaces with the weaker species. Thus, genetic drift may strengthen the stability of the range limits by weakening the stronger species. An extension of the model we used that includes genetic drift could test this hypothesis.

The evolutionary model we used allows for the evolution of trait mean and trait variance. As a result, species can evolve character displacement to partially release themselves from interspecific competition and coexist over a limited region of sympatry, before the swamping effects of asymmetric gene flow becomes sufficiently strong to halt their range expansion. However, the intrinsic competitive ability of the species determined by their characteristic parameters (e.g., R_*i*_, V_*i*_, or D_*i*_) was assumed to be fixed and did not evolve in our analyses. It is likely that in the real world species may evolve their competitive ability in response to interspecific competition [88]. Such species can experience different population outcomes as a consequence of this evolution [91, 92, 93]. If the evolution of competitive abilities is in the direction of releasing the species from interspecific competition, then no range limits may form by competition and the species may coexist all over the habitat. If competitive abilities are evolved to equalize the relative competitive strength of the species, without releasing them from competition, then the competitively formed range limits will eventually become (semi-) stable in the weak sense; noting that perfect equalization is unlikely to evolve. If the interface between the species reaches a sufficiently steep wiggle as they evolve to equalize their abilities, then their range limits gets strongly stabilized in the wiggle. Further, the steep gradient within the wiggle substantially shortens the overlap between the species (sharpens the range borders) and reduces the peripheral population density of the stronger species; see Figure 3. This may significantly slow down or halt further evolution of of the competitive ability of the species, since such abilities mainly evolve through overlapping populations. Therefore, we hypothesize that environmental wiggles not only can stabilize competitively formed range limits, but also they may halt evolution of species’ competitive ability by effectively isolating them from each other. The model we have used can be extended to include the evolution of competitive abilities in order to test our hypothesis.

## Appendix A: Model Equations

The equations of the model for two competing species, as well as a single solitary species are given Sections A.1 and A.2 below. The equations are the same as those developed in [24] with an additional term modeling Allee effects. The mathematical formulations given in [24] have some notational differences compared with the preceding foundational models [26, 20] which we preserve here. To facilitate comparisons with the preceding models, we provide a table of notational changes in Section A.4.

### A.1 The Two-Species Model

The final equations of the model for two competing species are presented by a system of partial differential equations that governs the evolution of each species’ population density *n*_*i*_(*x, t*), *i* = 1, 2, trait mean *q*_*i*_(*x, t*), and trait variance *v*_*i*_(*x, t*) at every habitat location *x* ∈ Ω and time *t* ∈ [0, *T*], *T >* 0. For brevity, let *u*:= (*n*_1_, *q*_1_, *v*_1_, *n*_2_, *q*_2_, *v*_2_) be a vector containing all model variables. Moreover, let the partial derivatives with respect to *t* and *x* be denoted by *∂*_*t*_ and *∂*_*x*_, respectively, and the second partial derivative with respect to *x* be denoted by 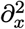. Then, the equation for the evolution of the population density *n*_*i*_ of the *i*th species is given by

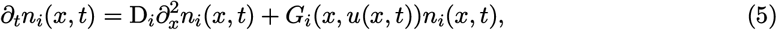

where *G*_*i*_(*x, u*) denotes the mean growth rate of the population as given by (8) below. Likewise, equations for the evolution of the trait mean *q*_*i*_ and the trait variance *v*_*i*_ within the *i*th species are given as

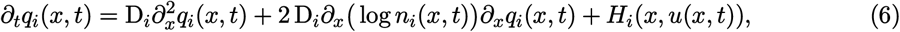

and

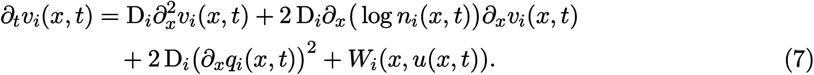

The nonlinear mappings *G*_*i*_, *H*_*i*_, and *W*_*i*_ used in (5)–(7) are defined as

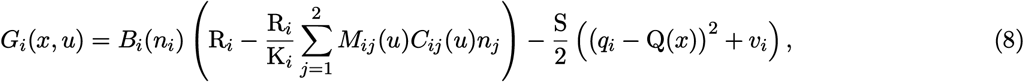

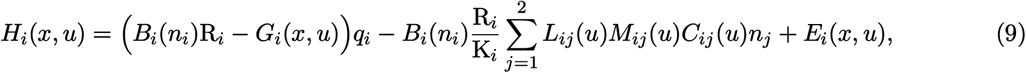

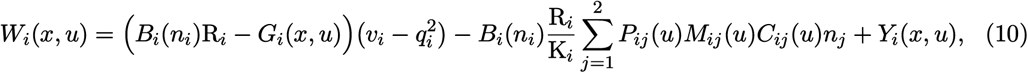

where, letting 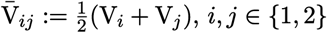,

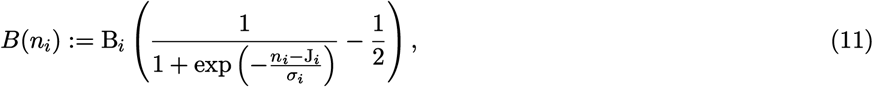

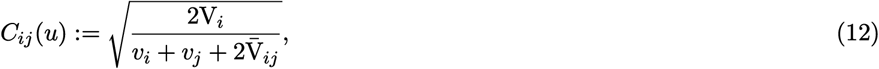

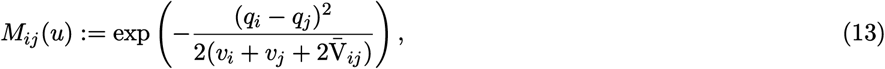

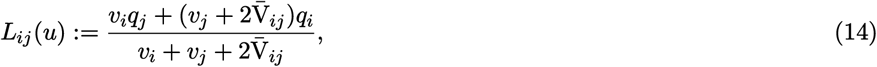

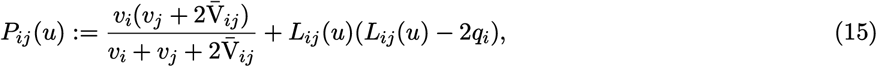

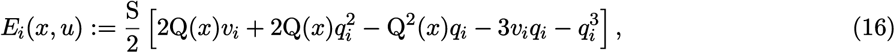

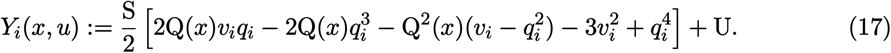

Table 1 of the main text gives the definitions of the model parameters and their default values used in the simulations.

Solving the equations of the model (5)–(7) over a habitat Ω = (*a, b*) requires specifying boundary conditions at habitat boundary points *x* = *a* and *x* = *b*. For the numerical studies presented throughout our work we always assume that there is no phenotype flux through the boundary of the habitat, which gives the homogeneous Neumann boundary conditions

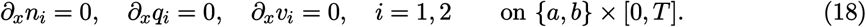

*∂*_*x*_*n*_*i*_ = 0, *∂*_*x*_*q*_*i*_ = 0, *∂*_*x*_*v*_*i*_ = 0, *i* = 1, 2 on {*a, b*} × [0, *T*]. (18) Equivalently, these conditions imply that the habitat boundary is reflecting.

#### A.2 The Single-Species Model

The equations of the model for a solitary species with symmetric intraspecific competition between phenotypes are given as

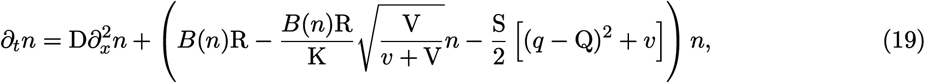

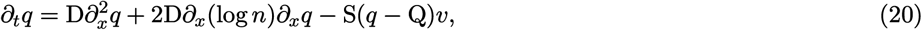

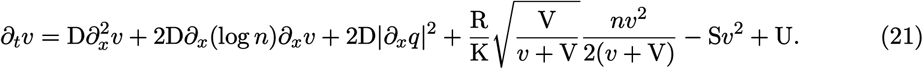

Since there exists only one species in the model, the numeration index *i* = 1 of the variables and parameters in (19)–(21) is dropped for notational simplicity. Further, the dependence of *n, q*, and *v* on *x* and *t*, as well as the dependence of Q on *x*, are not shown for the simplicity of exposition.

#### A.3 Assumption of Normal Distribution of Phenotypes

The model derivation assumption that requires the distribution of phenotypes to be normal is relatively strong and is made for the simplicity of model derivation in [24, 20, 26]. However, we do not expect the general results of our work to be significantly affected by the violation of this assumption, as long as the distribution of phenotypes remains to be approximately unimodal and symmetric (not heavily skewed) with finite moments. Within the framework of our study, such conditions are reasonably satisfied. In [20], the normality assumption is made following the assumptions that the trait is determined by many loci, allelic effects are additive within and between loci, individuals mate randomly, and selection is weak.

When the trait optimum changes linearly in space, gene flow is localized (e.g., diffusive), and the majority of genetic variation in the population is maintained by gene flow (migration) rather than by mutation, it is argued by Barton [25, 94] that continuum allelic effects at each loci can be reasonably assumed to be normal. If the allelic effects vary independently, then their sum (additive trait) will also be normally distributed. Knowing that the Central Limit Theorem extends (under certain conditions on the moments) to non-identically distributed random variables, the trait distribution will still tend to be normal if allelic effects are weakly dependent. The analysis performed by [24], as well as the single-species simulations we perform in the present work, confirm that the local phenotypic/genetic variation is determined predominantly by gene flow provided the species’ range evolves in sufficiently steep environmental gradients. This is a condition that holds in all of the simulations we perform in this work. Further, the nonlinearities that we study are in fact piecewise linear, with the nonlinear regions only occurring locally at the location of the knees. As we showed, these locations will be occupied by peripheral (low-density) populations of the species and do not contribute significantly to the gene flow across the core of the populations. Therefore, following Barton’s arguments, the normality assumption on the distribution of phenotypes should be fairly reasonable in our results. We also note that, the normality of allelic effects at each locus is indeed a much stronger condition than the normality of the overall distribution of phenotype values [94]. That is, the phenotypes can still be normally distributed even when the conditions we described above (high gene flow) is not satisfied.

#### A.4 Notational Differences with Preceding Models

In presenting the equations and the underlying components of the model, we used the same mathematical notations as in [24, 30]. To distinguish model variables from model parameters and mappings of model variables, to avoid conflicts with some commonly used notations in mathematical analysis of PDEs, and to allow for additional notational styles for possible future extensions and analyses of the model, some notational changes have been made in [24, 30], compared with the notations used in the foundational models of Kirkpatrick & Barton [26] and Case & Taper [20]. To facilitate comparisons with the preceding models, in Table 2 we provide the list of model parameters and variables, with the notations used in the present work (as well as in [24, 30]) and those used in [20] (as well as in [26] and other works in this family of models).

**Table 2.** Notational comparison between the model presented in this work and Case & Taper’s model [20]. The presence of the numeration index *i* in a variable implies that the variable can take a different value for each species.

| Parameter/variable/function description | Present work | Case & Taper’s [20] |
| --- | --- | --- |
| The random variable denoting phenotype values | $p$ | $z$ |
| Population density | $n_i$ | $N_i$ |
| Trait mean | $q_i$ | $\bar{z}_i$ |
| Trait variance | $v_i$ | $V_p$ |
| Per capita intrinsic growth rate | $g_i$ | $w_i$ |
| Mean intrinsic growth rate | $G_i$ | $\bar{w}_i$ |
| Competition kernel | $\alpha_{ij}$ | $\alpha$ |
| Relative frequency (distribution) of phenotypes | $\phi_i$ | $p_i$ |
| Mutational changes in frequency of phenotypes | $\partial_t^{(M)} \phi_i$ | — |
| Diffusion coefficient of species’ dispersal | $D_i$ | $D$ |
| Carrying capacity of the environment | $K_i$ | $K$ |
| Critical population density (Allee threshold) | $J_i$ | — |
| Maximum per capita growth rate | $R_i$ | $r_i$ |
| Variance of individuals’ phenotype utilization | $V_i$ | $V_u$ |
| Measure of the strength of natural selection | $S$ | $1/V_s$ |
| Rate of increase in trait variance due to mutation | $U$ | — |
| Optimal trait value for the environment | $Q$ | $\theta$ |

## Appendix B: Numerical Methods

In all simulation results presented in this work, the numerical solutions of the model equations are computed using the same numerical method as used in [24], with a slight modification regarding the terms *∂*_*x*_ log *n*_*i*_ as described below. The method is based on an implicit scheme with two stabilizing correction stages [95]. In each iteration of the scheme, instead of solving the nonlinear algebraic equations involved in the computations, the linearized version of these equations are solved. The iteration time steps are then made smaller to compensate for the linearization error. The first and second space derivatives are approximated using fourth-order centered differences. See Appendix B of the work by [24] for further details. In all simulations, we discretized the one-dimensional space (habitat) with a uniform mesh of size Δ*x* = 0.1 `X`. For the majority of the simulations in which we aimed to compute the equilibrium curves accurately, we used small time steps of length Δ*t* = 0.002 `T`. For simulations that required an exceedingly long evolutionary time horizon *T*, or did not require computation of fully accurate solutions, we chose longer time steps of Δ*t* = 0.005 `T`, Δ*t* = 0.01 `T`, or Δ*t* = 0.016 `T`.

The presence of the terms *∂*_*x*_ log *n*_*i*_ in the equations of the model, (6) and (7), make the numerical computations of the solutions particularly challenging. These terms are undefined when the population densities *n*_*i*_ are zero; see further details in [24, Section 4.1]. Even if we initialize the populations such that at least an infinitesimal population density is initially present everywhere, for example as *n*_*i*_ = sech(|*x*|), there is still a chance of reaching numerical singularities if the density of a species undergoes a long density decline (or extinction) regime, especially with our addition of the Allee effect to the model. To avoid this singularity problem, in our numerical computations we replace the terms *∂*_*x*_ log *n*_*i*_ with *∂*_*x*_ log(*n*_*i*_ + *ϵ*), *ϵ >* 0. A small value of *ϵ*, as small as 10^−5^, works well in our simulations and does not result in any noticeable difference in the solutions of the model compared with the original equations. With this minor modification, we could also initialize the population densities using bump functions, which are exactly equal to zero outside the support of the functions. We used the bump function

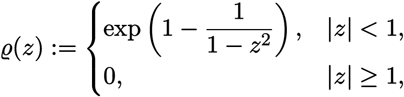

which has the compact support [−1, 1] and takes the maximum value 1 at *z* = 0.

## Supplementary Information

**Figure S1:**
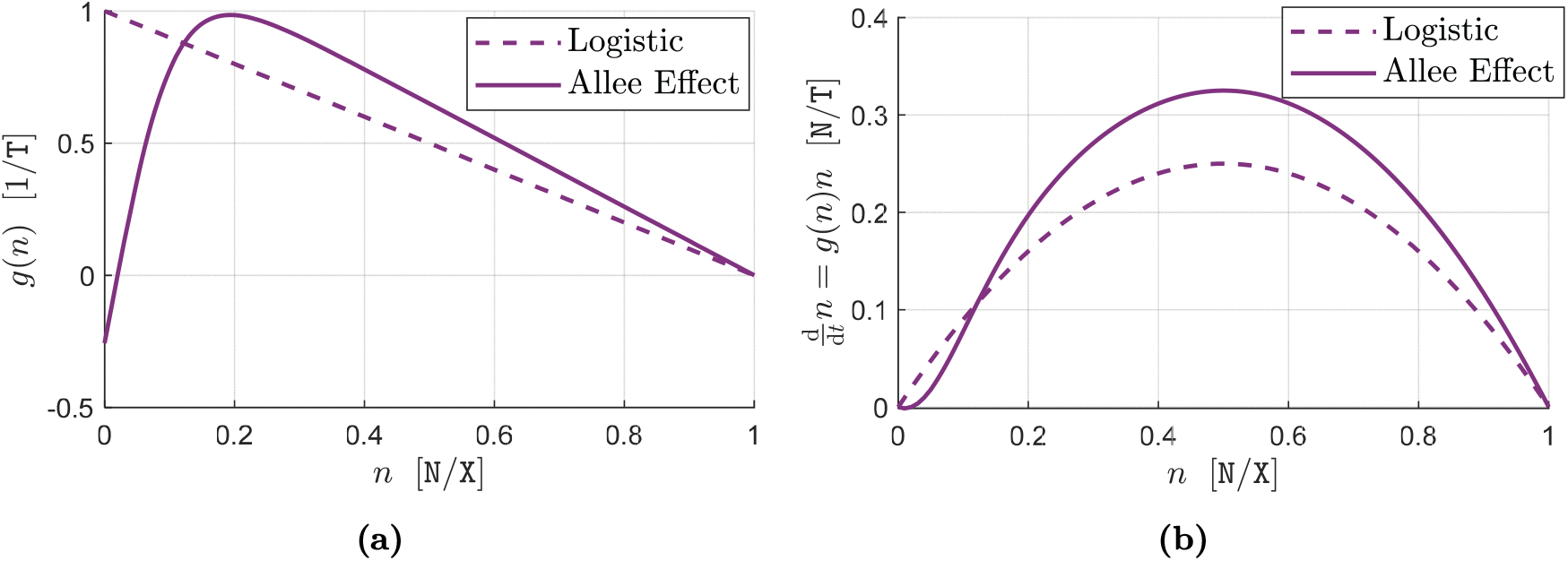
Intrinsic growth rate of a fully adapted population with Allee Effect. For a solitary (N = 1, hence we drop the numeration index *i*) population of individuals with phenotype *p*, which are completely generalist (V → ∞, which gives *α*(*p, p*′) → 1 for all *p* and *p*′), and are perfectly adapted to the environment (*p* = Q), the intrinsic growth rate given by equation (2) of the main text will depend only on the species’ population density, as *g*(*n*) = R (1 − *n/*K) *B*(*n*). In the absence of dispersal (a local isolated population) and mutations, the rate of change in the density of this population will be 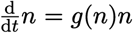. The graphs of *g*(*n*) and 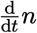 are shown in (a) and (b), respectively, both for the case where the growth rate is logistic, *B*(*n*) = 1, and the case where the population exhibits Allee Effect, 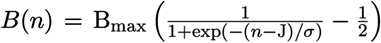. The parameter values used for computing the graphs are R = 1, K = 1, *J* = 0.02, B_max_ = 2.6, and = 0 05. Note that, as the curve with Allee Effect in (a) shows, for population densities below the critical density (*n <* J), the intrinsic growth rate *g* is negative.

**Figure S2:**
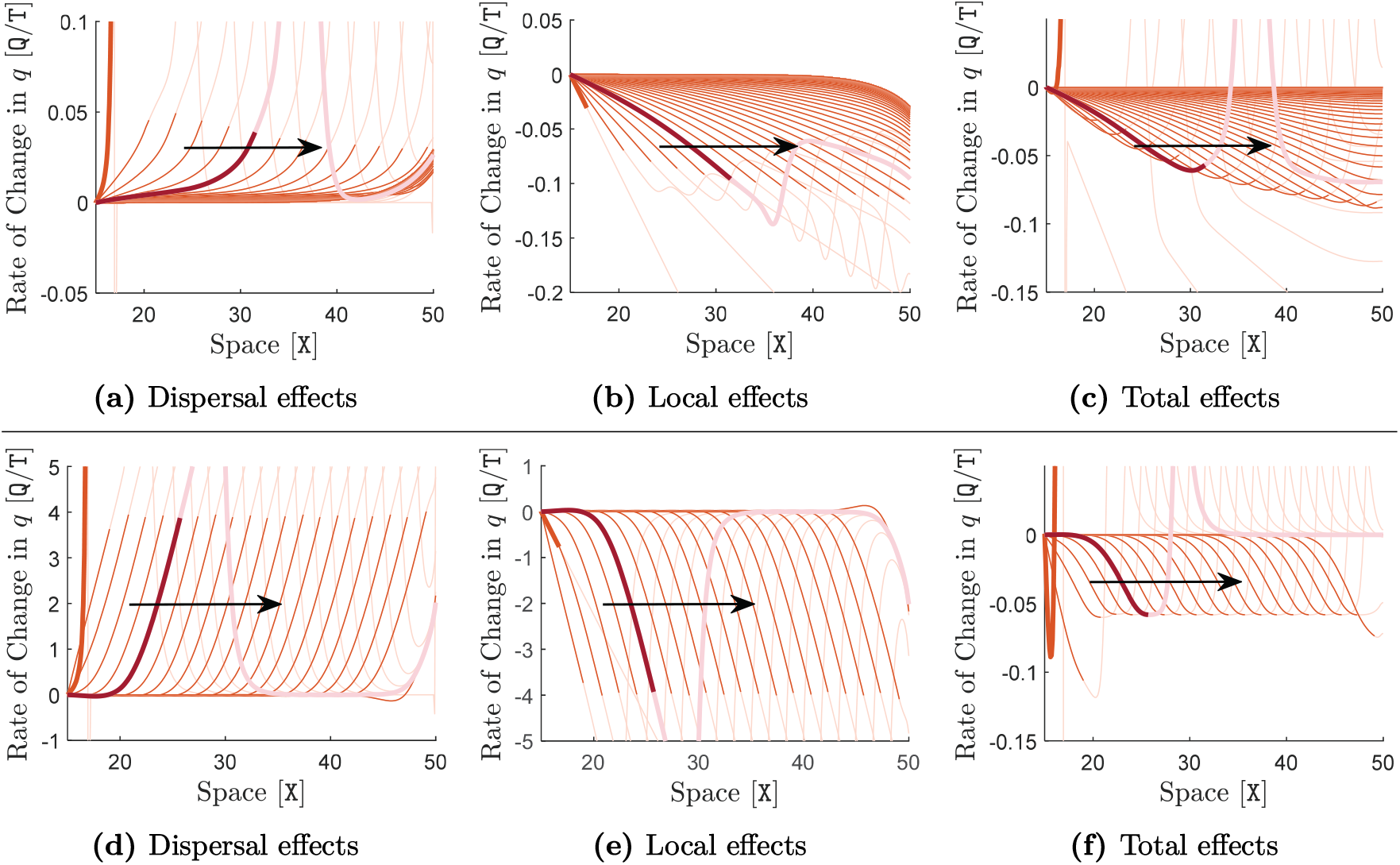
Effects of random gene flow and selection on population adaptation in shallow and steep environmental gradients. The same simulation as in Figure 1 of the main text is performed here, but with a shallow gradient of *∂*_*x*_Q = −0.1 `Q`/`X` for the graphs shown in the upper panel, and a steep gradient of *∂*_*x*_Q = −2.5 `Q`/`X` for the graphs shown in the lower panel. The contributions of dispersal (gene flow) to the rate of change of trait mean *∂*_*t*_*q* are shown in (a) and (d), the local contributions of selection and competitions are shown in (b) and (e), and total contribution are shown in (c) and (f). These contributions are computed in the same way as described in Figure 1 of the main text. The curves are shown only for the right-half of the species range (15 `X`, 50 `X`). The curves in the upper panel (shallow gradient) are shown at every 2 `T` for a simulation time horizon of *T* = 200 `T`, with a sample curve being highlighted in red at *t* = 10 `T`. The curves in the lower panel (steep gradient) are shown at every 40 `T` for a simulation time horizon of *T* = 800 `T`, with a sample curve being highlighted in red at *t* = 200 `T`. The same description as given in Figure 1 of the main text holds for the curve colors and arrows.

**Figure S3:**
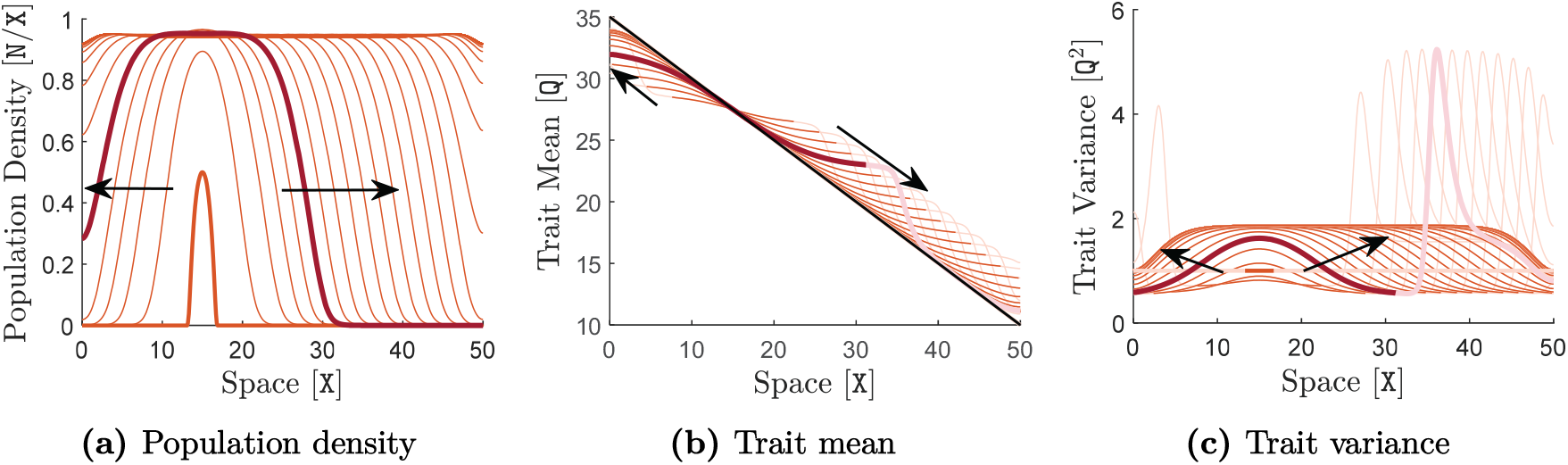
Range evolution of a solitary species with perfect initial adaptation. The same simulation as in Figure 1 of the main text is performed here, with the only difference being that the population is initialized to be perfectly adapted to the environment, that is, *q*(*x*, 0) = Q(*x*) for all *x* ∈ Ω. The same descriptions as given in Figures 1a–1c of the main text hold for the quantities shown in the graphs, curve colors, arrows, time increments between the curves, and the simulation time horizon.

**Figure S4:**
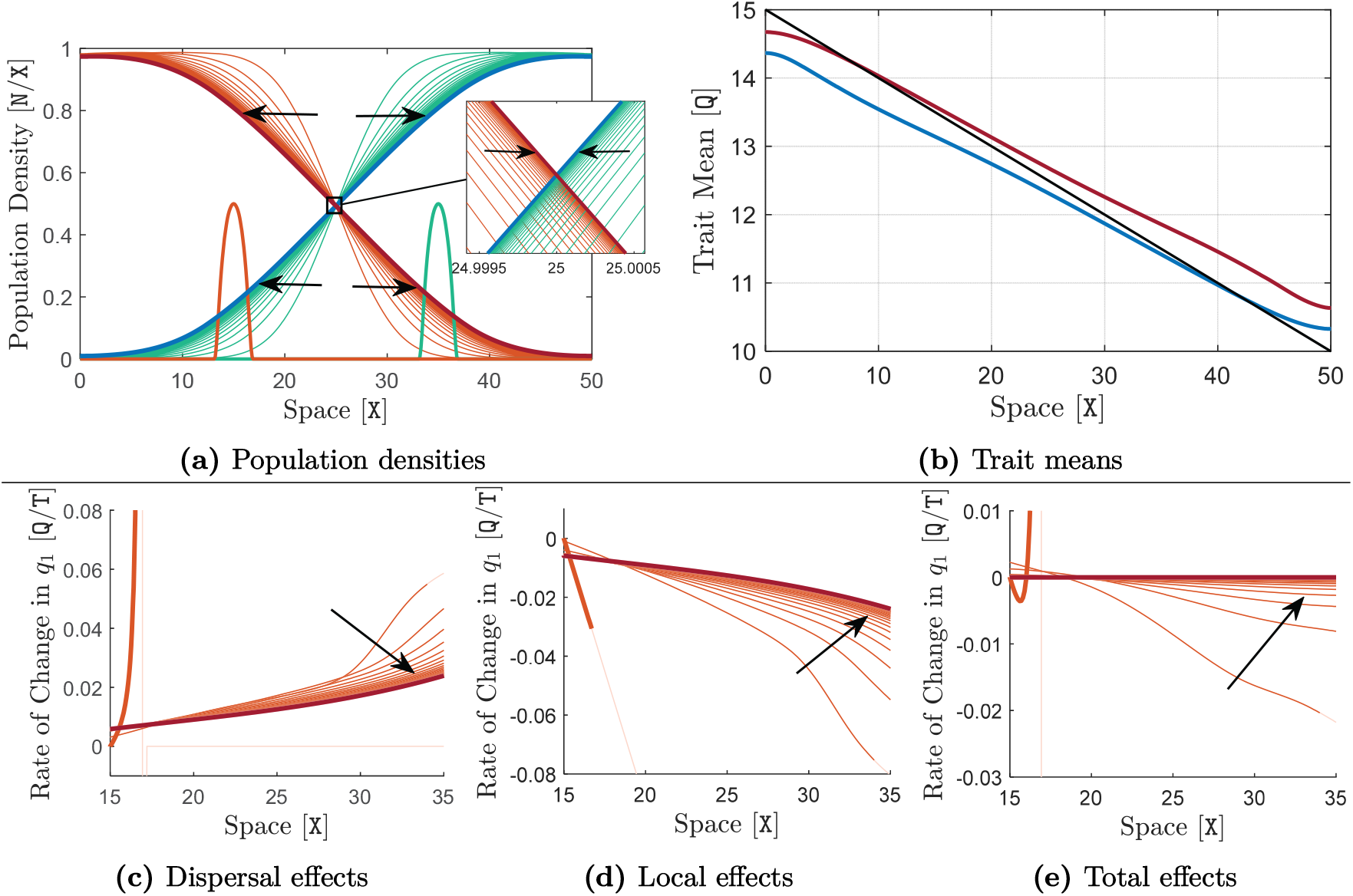
Evolution of character displacement and region of sympatric coexistence for two competitively identical species in a shallow environmental gradient. The same simulation as in Figure 2 of the main text is performed here, with the only difference being that the environmental gradient is set to be shallow. Specifically, the trait optimum Q is linear and decreasing, shown by the black line in (b), with a shallow gradient (slope) of *∂*_*x*_Q = −0.1 `Q`/`X`. The same descriptions as given in Figure 2 of the main text hold for the quantities shown in the graphs, curve colors, and arrows. Here, curves in (a) and (c)–(e) are shown at every 20 `T`, for a simulation time horizon of *T* = 1500 `T`. The initially formed region of sympatry in the middle of the habitat continues to expand, rather slowly, until the species become completely sympatric over the entire habitat.

**Figure S5:**
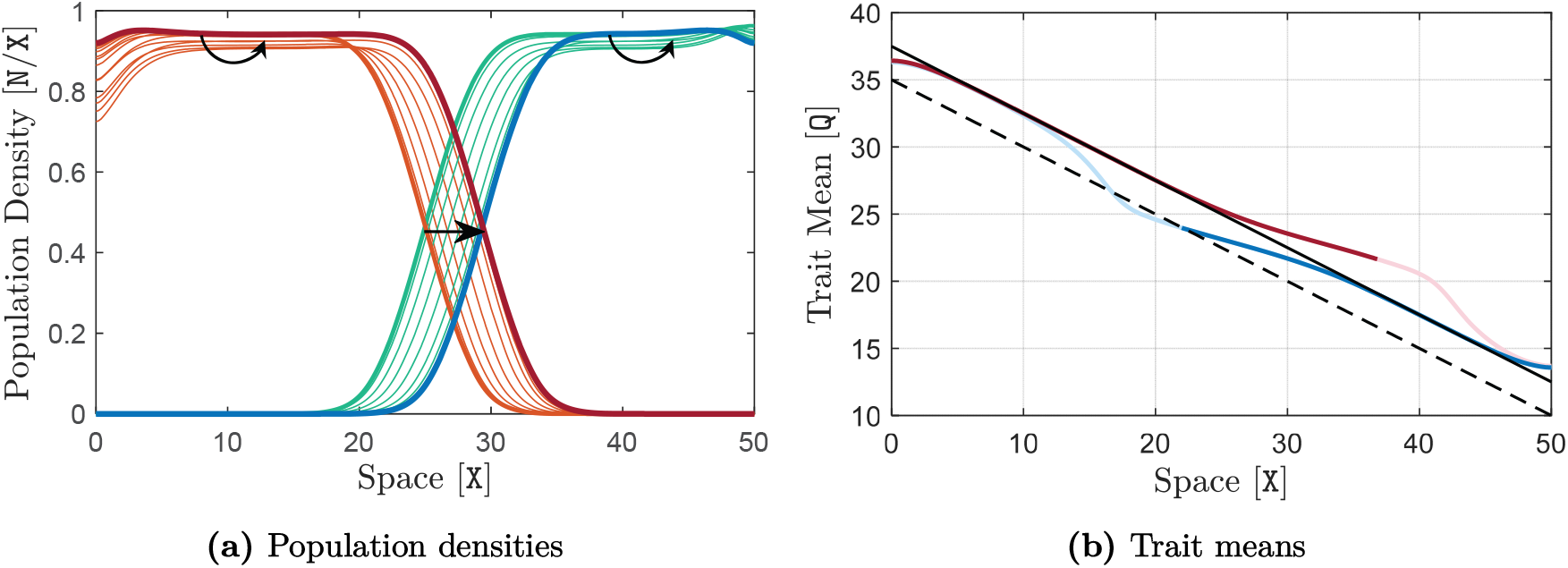
Permanent shift in the equilibrium region of sympatry and range limits in response to a permanent climate-warming disturbance. The two species are competitively identical, with the same parameters as used in Figure 2 in the main text. The species are initialized at *t* = 0 `T` by the final solution curves (equilibrium state) obtained at the end of the simulation in Figure 2 of the main text (with *∂*_*x*_Q = −0.5 `Q`/`X`). Starting from *t* = 0 `T`, the curve of trait optimum Q is gradually (linearly) shifted up by 2.5 Q over a time course of 10 `T`, and remains unchanged afterwards. That is, a climate-warming disturbance *δ*Q is added to the initial curve of Q, where *δ*Q = *αt/t*_r_ for 0 ≤ *t* ≤ *t*_r_ and *δ*Q = 0 for *t > t*_r_, with disturbance amplitude *α* = 2.5 Q and disturbance rise time *t*_r_ = 10 `T`; see Figure (*iv*) in Box 2 of the main text. The dashed black line in (b) shows the initial curve of Q, and the solid black line shows the completely shifted curve after *t* = 10 `T`. The simulation is performed for a time horizon of *T* = 500 `T`. The same description as given in Figure 2 of the main text holds for the curve colors and arrows. Curves of population density are shown at every 2 `T` in (a) and the final equilibrium curves obtained at *t* = 500 `T` are highlighted in red and blue. The final curves of trait mean obtained at *t* = 500 `T` are shown in (b). We observe that the range limits converge to a new equilibrium state after the climate-warming disturbance takes place, resulting in a permanent upslope shift in range limits and the region of sympatry.

**Figure S6:**
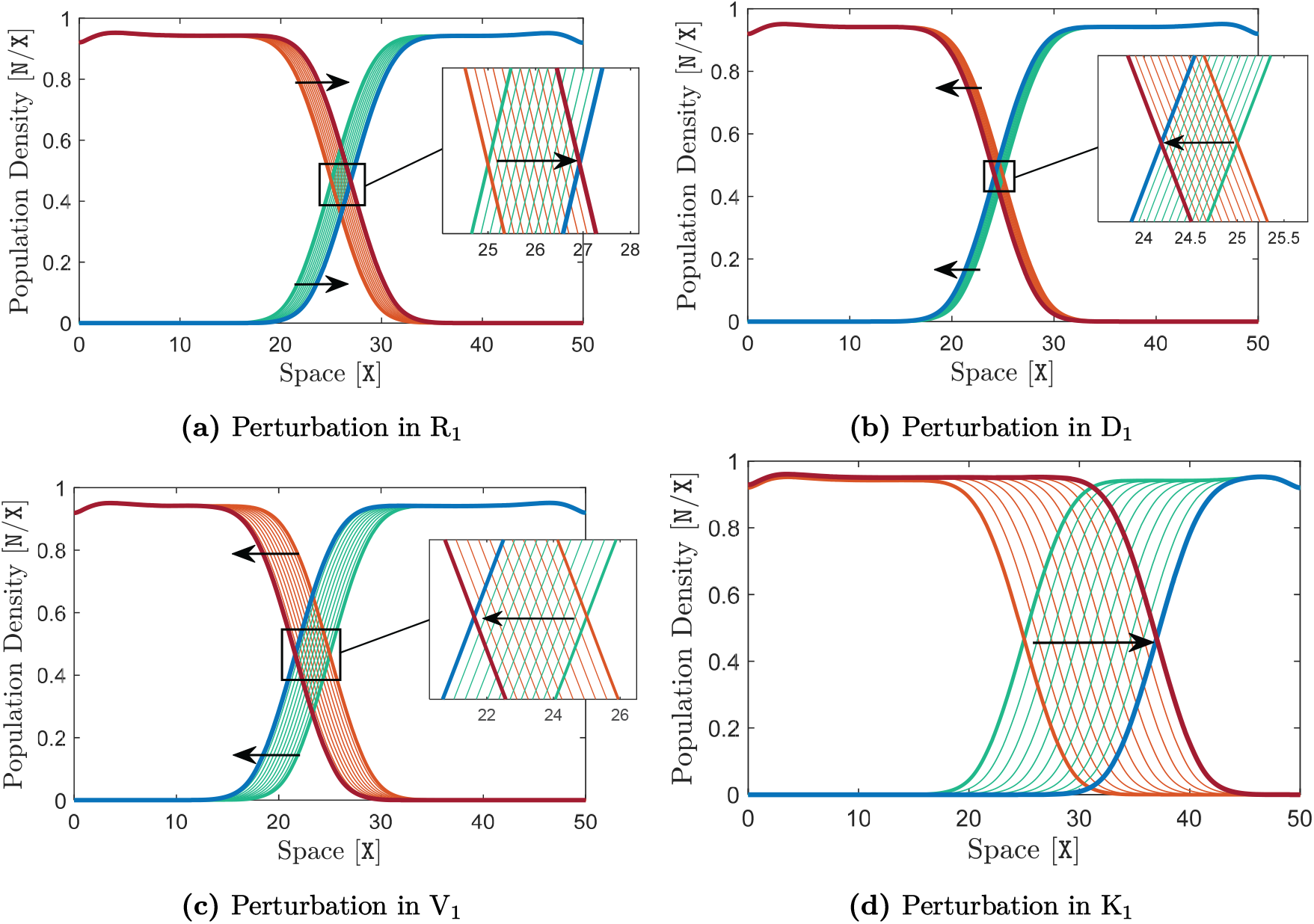
Instability of the range limits for competitively unequal species in a linear environment. The two species are initialized at *t* = 0 `T` by the final solution curves (equilibrium state) obtained at the end of the simulation in Figure 2 of the main text (with *∂*_*x*_Q = −0.5 `Q`/`X`). Both species have the same parameter values, equal to those used in Figure 2, except for one of the parameters that is increased by only one percent for the downslope species in each panel. In (a), R_1_ is increased to R_1_ = 1.01 `T` at the beginning of the simulation and is kept constant at this value for the entire simulation. This makes the downslope species slightly stronger than the upslope species. Similarly, in (b), D_1_ is increased by one percent to D_1_ = 1.01 `X`^2^*/*`T`, making the downslope species slightly weaker. In (c), V_1_ is increased to V_1_ = 9.09 `Q`^2^, which makes the downslope species weaker. In (d), K_1_ is increased to K_1_ = 1.01 N*/*X, making the downslope species stronger. In each case, the simulation is performed for a time horizon of *T* = 1000 `T`. Curves of population density are shown at every 100 `T` in each graph and the final curves at *t* = 1000 `T` are highlighted. The same description as given in Figure 2 of the main text holds for the curve colors and arrows. In all cases, we observe that a slight change in the competitive equivalence of the two species destabilizes (eliminates) the initial equilibrium (formed for the non-generic case of identical species), causing a constant shift in the range limits towards the weaker species.

**Figure S7:**
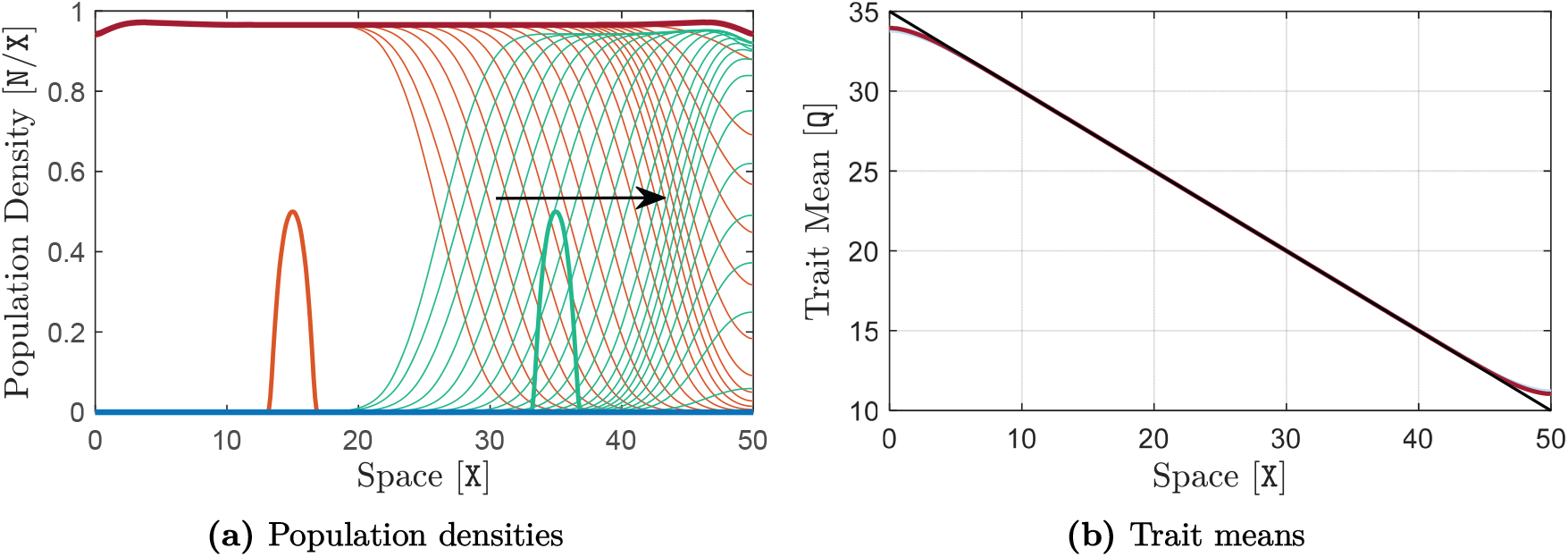
Competitive exclusion. The downslope species is significantly stronger, with a maximum growth rate of R_1_ = 1.2 `T`^−1^ versus the growth rate R_2_ = 1 `T`^−1^ of the upslope species. The rest of the model parameters for both species are the same and equal to the values given in Table 1 of the main text. The curves of population density of the two species are shown in (a) as their range evolves in time. Arrows show the direction of evolution in time after the region of sympatry is formed between the species and starts moving upslope. The same description as given in Figure SS4 holds for curve colors, and the highlighted curves in red and blue are the final curves obtained at the end of simulations. In (a), the curves are shown at every 40 `T` for a simulation time horizon of *T* = 1500 `T`. The downslope species eventually excludes the upslope species from the habitat. The curves of steady-state trait means obtained at the end of the simulation are shown in (b).

**Figure S8:**
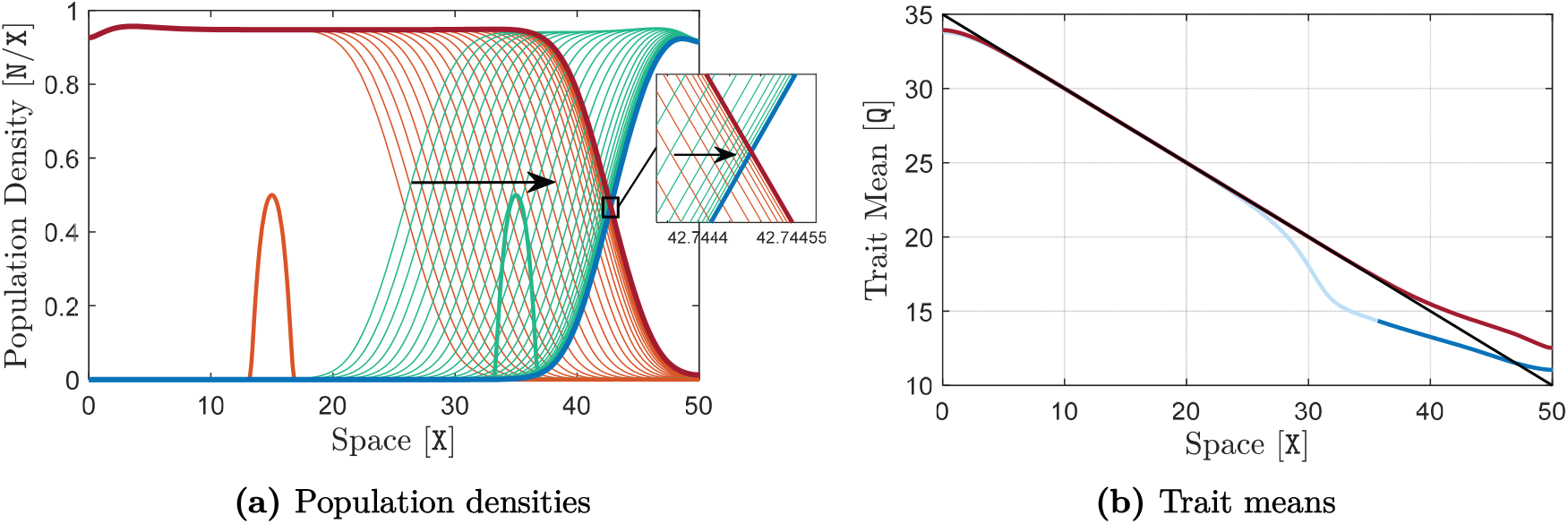
Marginal coexistence. The downslope species is slightly stronger, with a maximum growth rate of R_1_ = 1.05 `T`^−1^ versus the growth rate R_2_ = 1 `T`^−1^ of the upslope species. The rest of the model parameters for both species are the same and equal to the values given in Table 1 of the main text. The curves of population density of the two species are shown in (a) as their range evolves in time. Arrows show the direction of evolution in time after the region of sympatry is formed between the species and starts moving upslope. The same description as given in Figure SS4 holds for curve colors, and the highlighted curves in red and blue are the final curves obtained at the end of simulations. In (a), the curves are shown at every 100 `T` for a simulation time horizon of *T* = 6000 `T`. The species’ population distribution converges to an evolutionarily stable equilibrium state, at which the upslope species survives marginally at the vicinity of the habitat boundary. The curves of steady-state trait means obtained at the end of the simulation are shown in (b).

**Figure S9:**
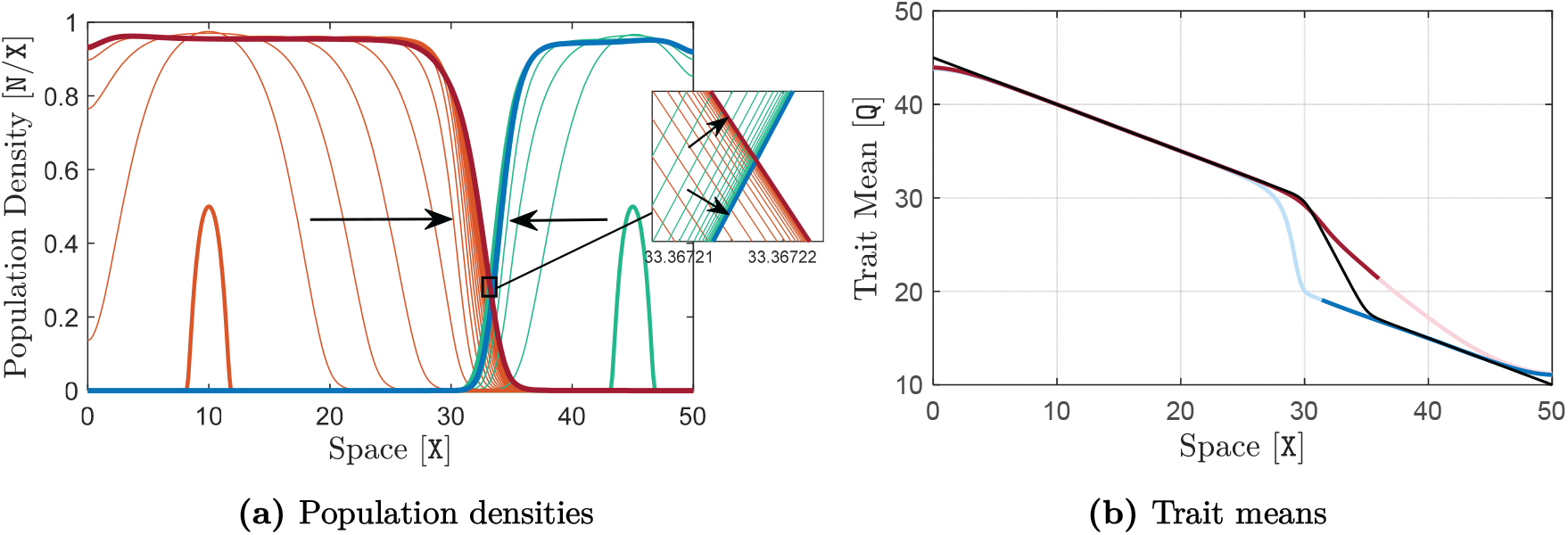
Formation of stable range limits at an environmental wiggle when the two species are initialized at opposite sides of the wiggle. A similar simulation to that of the lower panel of Figure 3 in the main text is performed here, with the main difference being that the populations are initialized at the opposite sides of the wiggle. That is, R_1_ = 1.1 `T`^−1^, R_2_ = 1 `T`^−1^, and the rest of the parameters take their typical values given in Table 1 of the main text. The habitat has an environmental wiggle. The slope of the trait optimum switches sharply from −0.5 `Q`/`X` to −2.5 `Q`/`X` between the two knees of the wiggle located at *x* = 30 `X` and *x* = 35 `X`. The simulation is performed for a time horizon of *T* = 750 `T` and curves of population density are shown in (a) at every 8 `T`. The same description as given in Figure 2 of the main text holds for the curve colors and arrows. The curves highlighted in red and blue in (a) are associated with the equilibrium state of the populations, reached (approximately) at the end of the simulations (*t* = 750 `T`). These curves represent the evolutionarily stable range limits formed at the environmental wiggle. The corresponding curves of equilibrium trait mean are shown in (b), where the black line shows the environmental trait optimum Q. The equilibrium curves of trait mean are made transparent over the regions where population densities are approximately zero.

**Figure S10:**
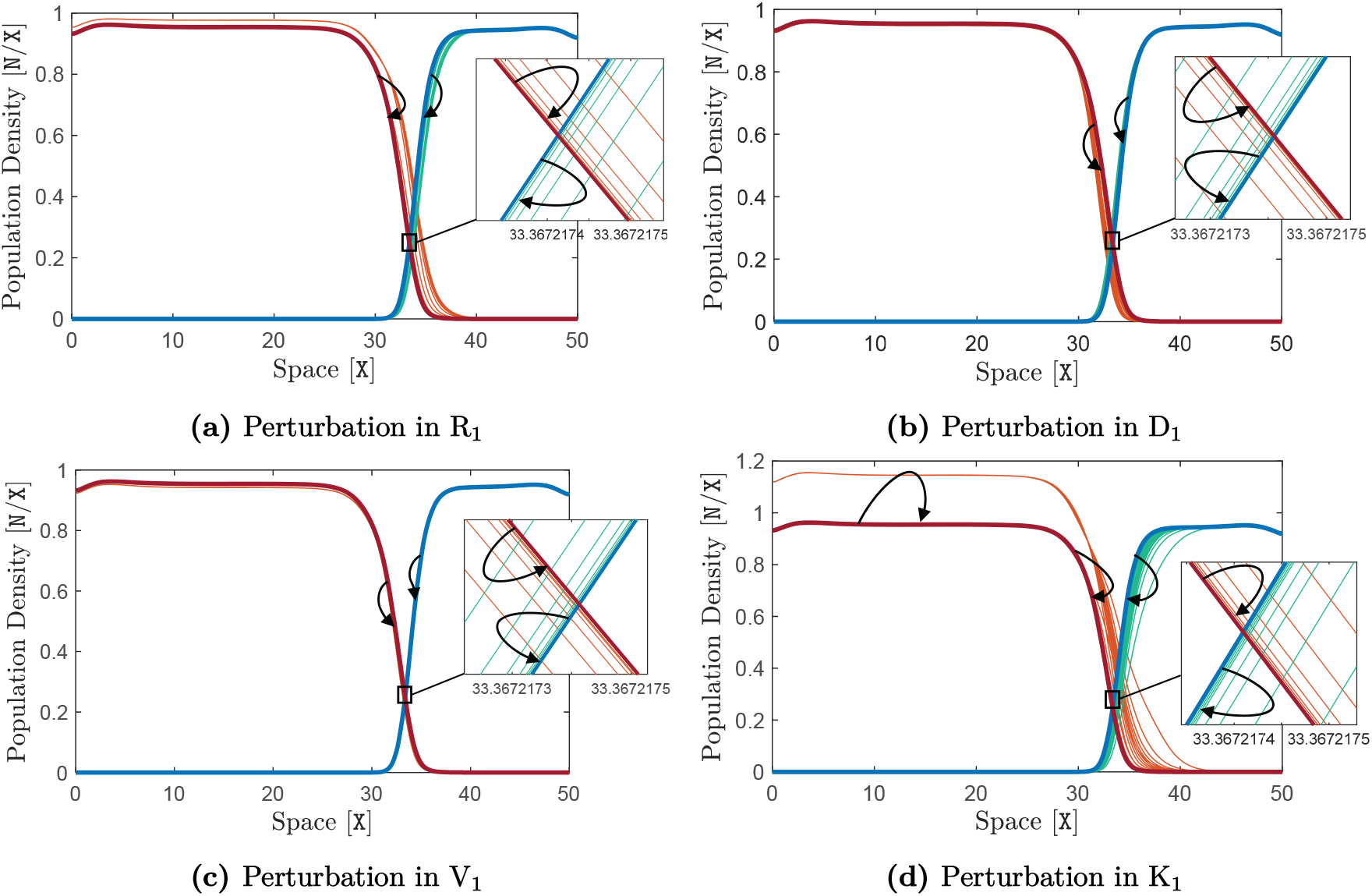
Robustness of the stability of the range limits formed at wiggles against transient perturbations in species’ parameters. The two species are initialized at *t* = 0 `T` by the final solution curves (equilibrium state) obtained at the end of the simulation shown in the lower panel of Figure 3 in the main text (in which the range limits are stabilized at an environmental wiggle). Both species take the same parameter values as used in Figure 3, except for one of the parameters that is perturbed for the downslope species in each panel by an additive rectangular-shaped perturbation of amplitude *α* = 20% and duration *t*_e_ = 200 `T`; see Figure (*iii*) in Box 2 of the main text. In (a), R_1_ is increased by 20% to the value R_1_ = 1.32 `T` at the beginning of the simulation, is kept constant at this value for a period of 200 `T`, and then is decreased back to its initial value R_1_ = 1.1 `T`. Similarly, in (b), D_1_ is increased by 20% to D_1_ = 1.20 `X`^2^*/*`T`, and then is decreased back to its initial value D_1_ = 1 `X`^2^*/*`T` after 200 `T`. In (c), V_1_ is perturbed by 20% over a time period of 200 `T`. In (d), a similar perturbation is applied to K_1_. In each case, the simulation is performed for a time horizon of *T* = 1500 `T`. Curves of population density are shown in each graph at every 20 `T`, and the final curves at *t* = 1500 `T` are highlighted. The same description as given in Figure 2 of the main text holds for the curve colors and arrows. In all cases, we observe that the stability of the range limits formed at the wiggle is maintained under moderately strong transient perturbations that make the downslope (stronger) species temporarily stronger (in (a) and (d)) or weaker (in (b) and (c)). In particular, the initial equilibrium range limits are precisely restored at the end of the simulation, as the perturbations only occur transiently for a duration of 200 `T` at the beginning of the simulation.

**Figure S11:**
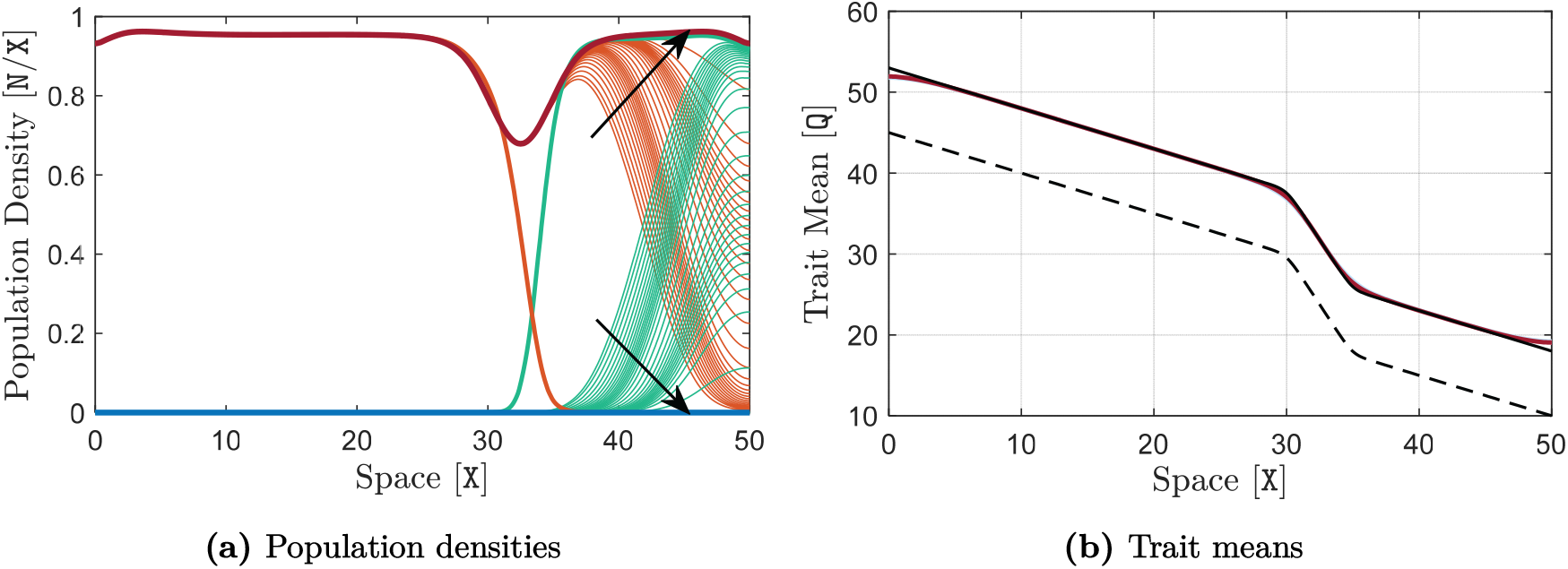
Destabilization of the range limits by a strong climate-warming disturbance when the downslope species is stronger, leading to the exclusion of the upslope species. The same simulation as presented in Figure 6 of the main text is further continued here until a longer evolution time of *T* = 2000 `T`. Curves of population density are shown in (a) at every 50 `T`. The final curves at *t* = 2000 `T` are highlighted in red and blue, and their corresponding trait mean curves are shown in (b). The same description as given in Figure 5 of the main text holds for curve colors, the arrow, and the dashed curve. Note that the final curve of trait mean for the upslope species is made entirely transparent since the final population density of this species is equal to zero everywhere in the habitat and hence its trait mean is not biologically meaningful.

**Figure S12:**
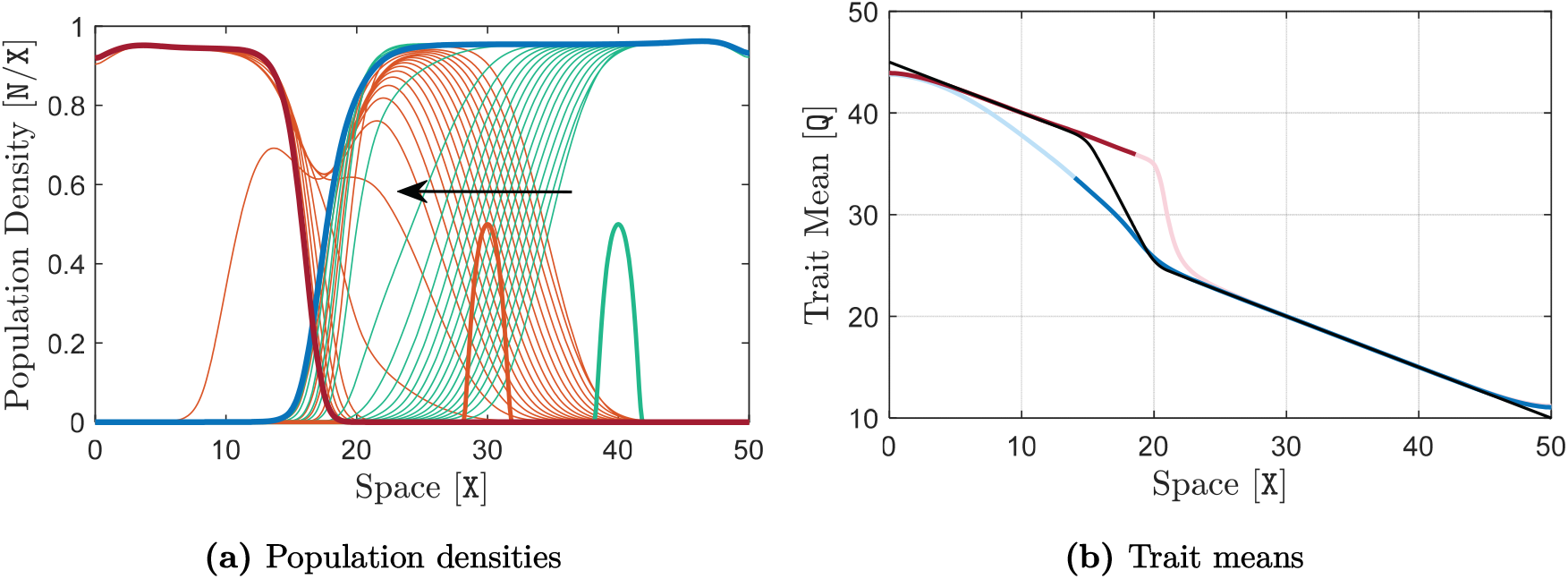
Formation of stable range limits at an environmental wiggle when the upslope species is stronger. A similar simulation to the one presented in the lower panel of Figure 3 in the main text is performed here, with the main difference being that here the upslope species is made stronger. That is, we set R_1_ = 1 `T`^−1^ and R_2_ = 1.1 `T`^−1^. The rest of the parameters take their typical values given in Table 1 of the main text. The simulation is performed for a time horizon of *T* = 1500 `T` and curves of population density are shown in (a) at every 30 `T`. The same description as given in Figure 2 of the main text holds for the curve colors and the arrow. The curves highlighted in red and blue are associated with the equilibrium state of the populations, reached (approximately) at the end of the simulation (*t* = 1500 `T`). The highlighted equilibrium curves in (a) represent the evolutionarily stable range limits formed at the environmental wiggle, with their corresponding curves of trait mean shown in (b). The black line in (b) shows the environmental trait optimum Q, whose slope switches sharply from −0.5 `Q`/`X` to −2.5 `Q`/`X` between the two knees of the wiggle located at *x* = 15 X and *x* = 20 X. The curves of trait mean in (b) are made transparent over the regions where population densities are approximately zero, as the values of trait mean over these regions are not biologically meaningful.

**Figure S13:**
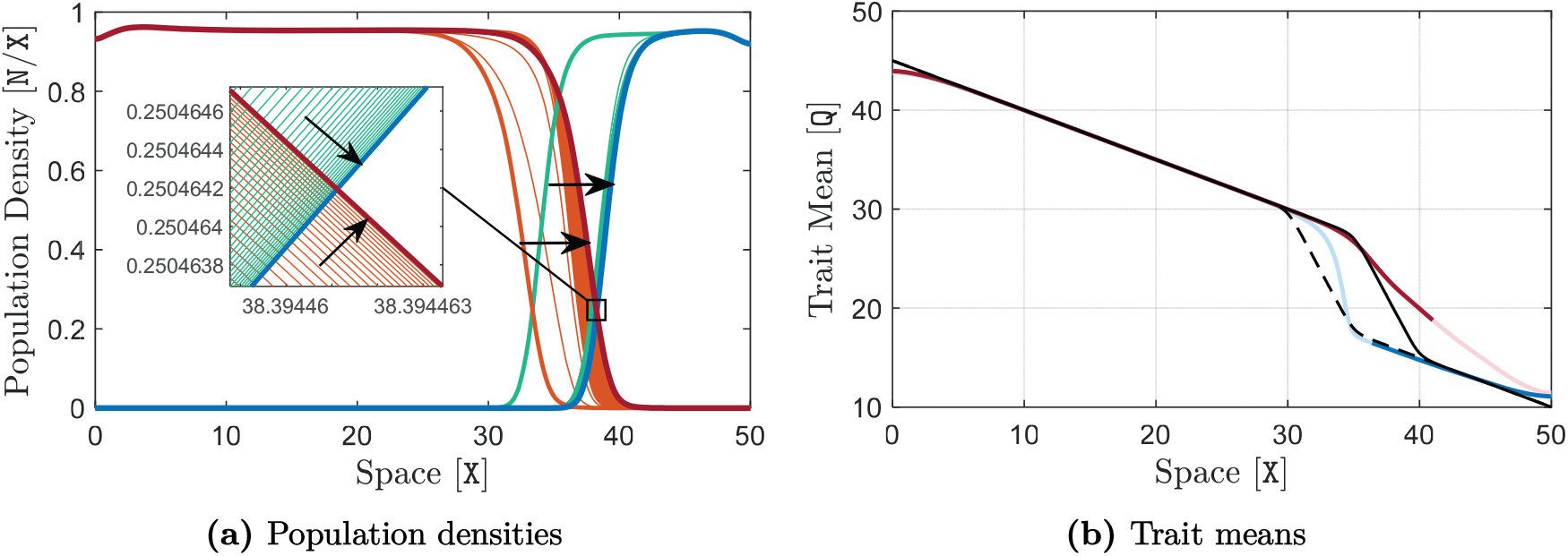
Range shifts resulting from an environmental disturbance that moves the wiggle towards the weaker species. The simulation is performed using the same model parameters as used in Figure 3 of the main text. The equilibrium curves obtained at the end (*t* = 1500 `T`) of the simulation shown in Figure 3 of the main text are used as the initial curves here. At *t* = 0 `T`, the wiggle in the trait optimum Q is immediately moved upslope by 5 `X` and remains unchanged afterwards. The initial curve of Q is shown by the dashed black line in (b), and the shifted curve after *t* = 0 `T` is shown by the solid black line. The simulation is performed for a time horizon of *T* = 1500 `T`. Curves of population density are shown at every 2 `T` in (a), and the final curves obtained at *t* = 1500 `T` are highlighted. The final curves of trait mean obtained at *t* = 1500 `T` are shown in (b). The same description as given in Figure 2 of the main text holds for the curve colors. We observe that the range limits move and get stabilized at the new location of the wiggle.

**Figure S14:**
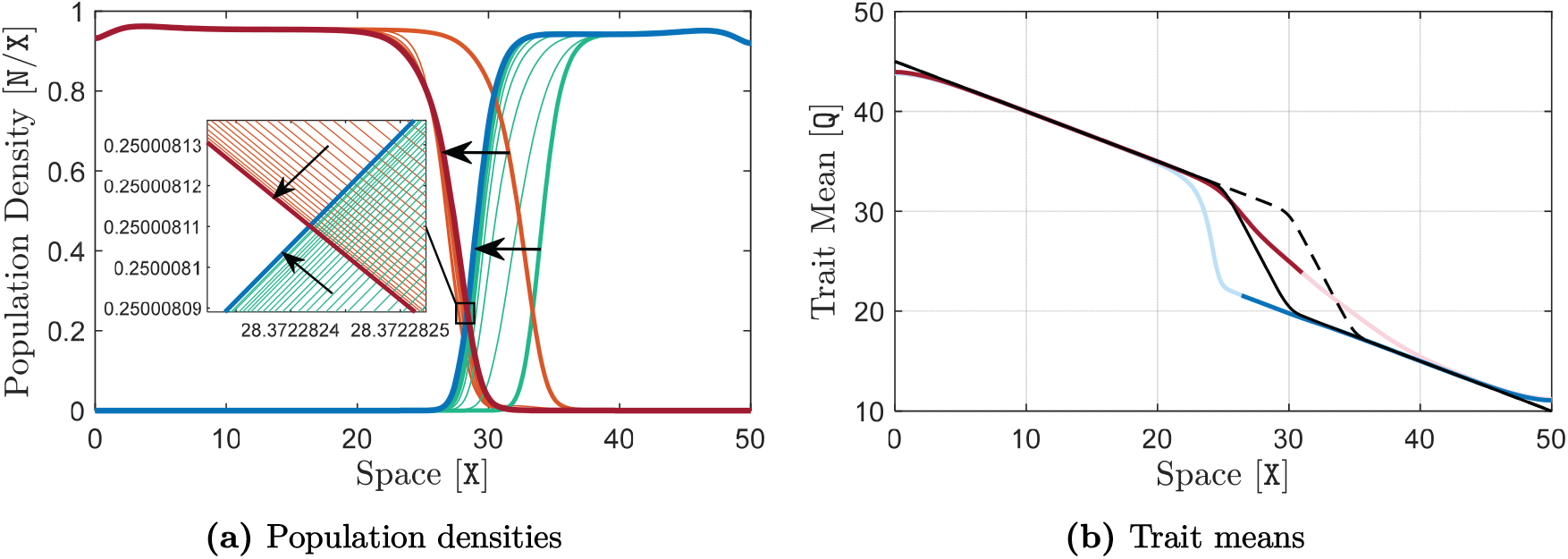
Range shifts resulting from an environmental disturbance that moves the wiggle towards the stronger species. The same simulation as in Figure SS13 is performed here, but with a disturbance that moves the wiggle downslope by 5 `X`. Since the disturbance is sufficiently small (even though it is a relatively large disturbance), we observe that the range limits move and get stabilized at the new location of the wiggle.

